# Perfluorooctanoic Acid Induces Liver and Serum Dyslipidemia in Humanized PPARα Mice fed an American Diet

**DOI:** 10.1101/2021.04.03.438316

**Authors:** JJ Schlezinger, T Hyötyläinen, T Sinioja, C Boston, H Puckett, J Oliver, W Heiger-Bernays, TF Webster

## Abstract

Per- and polyfluoroalkyl substances (PFAS) are pervasive in the environment resulting in nearly universal detection in people. Human serum PFAS concentrations are strongly associated with increased serum low-density lipoprotein cholesterol (LDL-C), and growing evidence suggests an association with serum triacylglycerides (TG). Here, we tested the hypothesis that perfluorooctanoic acid (PFOA) dysregulates liver and serum triacylglycerides in human peroxisome proliferator activated receptor α (hPPARα)-expressing mice fed an American diet. Mice were exposed to PFOA (3.3 mg/l) in drinking water for 6 weeks resulting in a serum concentration of 48 ± 9 μg/ml. In male and female hPPARα mice, PFOA increased total liver TG and TG substituted with saturated and monounsaturated fatty acids. Lack of expression of hPPARα alone also increased total liver TG, and PFOA treatment had little effect on liver TG in null mice. In hPPARα mice, PFOA neither significantly increased nor decreased serum TG; however, there was a modest increase in TG associated with very low-density cholesterol particles in both sexes. Across studies, a non-monotonic effect of PFOA on serum TG is evident, with the serum PFOA concentration in this study falling near the “null point” between increasing and decreasing serum TG. Intriguingly, in female PPARα null mice, PFOA significantly increased serum TG, with a similar trend in males. PFOA also modified fatty acid and TG homeostasis-related gene expression in liver, in a hPPARα-dependent manner, but not in adipose. The results reveal the importance of context (serum concentration and genotype) in determining the effect of PFOA on lipid homeostasis.

## Introduction

Per- and polyfluoroalkyl substances (PFAS) are extensively used in fluoropolymer production, consumer products (i.e., stain- and water-repellent fabrics, food contact materials, polishes, waxes, paints) and aqueous fire-fighting foam. At least 50% of public drinking water samples tested in the US contained two or more PFAS (Guelfo et al., 2018), exposing 10 million Americans or more to concentrations of PFAS that exceed guidelines set by multiple regulatory authorities (Hu et al., 2016). Daily exposure to PFAS occur via contaminated food, drinking water, personal care products, dust and air (De Silva et al., 2021; EFSA, 2018), resulting in nearly universal detection in the US population (CDC, 2019). Many PFAS are extremely persistent and mobile in the environment and have long half-lives (years) in humans (Cousins et al., 2016; Scheringer et al., 2014).

A number of adverse health endpoints are associated with PFAS exposure as observed in epidemiological studies including birth outcomes, immunologic effects, and metabolic disruption. Increased concentrations of serum total cholesterol, non-high density lipoprotein cholesterol (HDL-C), and low density lipoprotein cholesterol (LDL-C) are among the best supported, most sensitive endpoints in both cross-sectional and longitudinal epidemiology studies (reviewed in (ATSDR, 2018; EFSA, 2018) and recent studies (Dong et al., 2019; Graber et al., 2019; He et al., 2018; Jain and Ducatman, 2019; Li et al., 2020; C.-Y. Lin et al., 2020; Liu et al., 2020; Seo et al., 2018)). Increased serum triacylglycerides is also a sensitive effect, but not as well supported by epidemiology studies (Canova et al., 2020; Gardener et al., 2020; Jain and Ducatman, 2019; C. Y. Lin et al., 2020; Seo et al., 2018; Steenland et al., 2010, 2009; Zeng et al., 2015). Fewer studies have reported null or negative associations (T.-W. Lin et al., 2020; Matilla-Santander et al., 2017).

Conflicting results on the relationship between PFAS and serum triglyceride homeostasis have been generated in rodent models. In mice fed a standard, low fat/low cholesterol rodent diet and treated with high perfluorooctanoic acid (PFOA) doses show decreased serum triglyceride and total cholesterol levels (reviewed in (Pouwer et al., 2019; Rebholz et al., 2016)), with opposite effects observed in the epidemiology. But, some studies do indicate treatment of mice with moderate doses of PFOA induces hypercholesterolemia and hypertriacylglyceridemia when mice are fed a cholesterol and fat containing diet does (Rebholz et al., 2016; Yan et al., 2014).

A second conundrum is that PFAS share a potential molecular initiating event (MIE), namely activation of peroxisome proliferator activated receptor α (PPARα), with fibrate drugs, which are used to reduce blood lipids (Fruchart and Duriez, 2006; Kersten and Stienstra, 2017; Staels et al., 1998), identifying this MIE as critical to two seemingly divergent pathways. Studies using reporter assays demonstrate that PFAS of multiple chain lengths, both carboxylated and sulfonated, activate mouse and human PPARα (hPPARα) (Maloney and Waxman, 1999; Rosenmai et al., 2018; Takacs and Abbott, 2007; Vanden Heuvel et al., 2006). Multiple studies have shown that hPPARα-dependent gene expression is induced by PFAS in human hepatocyte models, including primary hepatocytes (Bjork et al., 2011; Buhrke et al., 2015, 2013; Peng et al., 2013; Rosenwald et al., 2017; Wolf et al., 2012, 2008). However, PPARα was found to account for only 80-90% of PFAS regulated genes in comparisons between PPARα wildtype and knockout mice (Rosen et al., 2017), indicating that PFAS initiate multiple MIEs.

A third issue is that there are species differences in ligand specificity between PPARα in mice and humans. Differences in the ligand binding domains between mouse and human PPARα are thought to contribute to differences in ligand specificity and gene expression patterns (Keller et al., 1997; Oswal et al., 2014; Rakhshandehroo et al., 2009). In particular, activation of PPARα in mice results in peroxisome proliferation and dysregulation of cell cycle genes, which does not occur in humans (Morimura et al., 2006).

Studies of effects of PFAS on liver and serum lipids and their lipidomes have largely been conducted in rodent models. PFOA-induced hepatosteatosis has been observed previously in mice expressing wildtype PPARα (mPPARα), human PPARa (hPPARα) mice and PPARα null mice (Das et al., 2017; Minata et al., 2010; Nakagawa et al., 2012; T. Nakamura et al., 2009; Tan et al., 2013). While hepatosteatosis is induced in PFOA-exposed mice fed standard rodent diets, the severity of the hepatosteatosis is increased when mice are co-exposed to PFOA and a high fat diet (Tan et al., 2013). Importantly, in an exposure scenario that generated an approximately steady state PFOA body burden, hPPARα mice were more susceptible to hepatic steatosis than mPPARα mice (Nakagawa et al., 2012). Effects of PFAS on serum triacylglycerides and the serum lipidome have only been conducted in rodent models, and as noted above, the effects are dependent upon exposure level and diet (reviewed in (Pouwer et al., 2019; Rebholz et al., 2016),(Pfohl et al., 2020; Yan et al., 2014)).

Here, we tested the hypothesis that PFOA exposure induces liver and serum hypertriacylglyceridemia in mice expressing hPPARα. Further, we compared the results to those in PPARα null mice to examine the contribution of PPARα. Both sexes were studied. Adipose tissue is also a significant source of fatty acids, and thus, contributor to triacylglyceride homeostasis in liver (Alves-Bezerra and Cohen, 2017). Thus, expression of genes involved in fatty acid and triacylglyceride homeostasis in liver and adipose were quantified. The data document modulation of multiple triacylglyceride types by PFOA exposure in liver and serum, only some of which were dependent on PPARα, along with important sex differences in a human relevant model.

## Materials and Methods

### Materials

Perfluorooctanic acid (171468, 95% pure) was from Sigma-Aldrich (St. Louis, MO). All other reagents were from Thermo Fisher Scientific (Waltham, MA), unless noted.

### In vivo exposure

All animal studies were approved by the Institutional Animal Care and Use Committee at Boston University and performed in an American Association for the Accreditation of Laboratory Animal Care accredited facility (Animal Welfare Assurance Number: A3316-01). Data from these mice have been previously reported (Schlezinger et al., 2020). Male and female, humanized PPARα mice (hPPARα) were generated from mouse PPARα null, human PPARα heterozygous breeding pairs (generously provided by Dr. Frank Gonzalez, NCI)(Yang et al., 2008). Experiments were carried out using 11 cohorts of mice generated from four breeding pairs (**Table S1**). Genotyping for mouse and human PPARα was carried out by Transnetyx (Cordova, TN). The expression level of hPPARα in liver was confirmed by RT-qPCR.

At weaning, mice were provided a custom diet based on the “What we eat in America (NHANES 2013/2014)” analysis for what 2-19 year old children and adolescents eat (Research Diets, New Brunswick, NJ)(USDA, 2018). The diet contains 51.8% carbohydrate, 33.5% fat, and 14.7% protein, as a % energy intake (**Table S2**). Fats are in the form of soybean oil, lard and butter, with cholesterol at 224 mg/1884 kcal. Vehicle (Vh) and treatment water were prepared from NERL High Purity water (23-249-589, Thermo Fisher Scientific), which is prepared using the most efficacious methods to remove PFAS (i.e., reverse osmosis and carbon filtering)(Appleman et al., 2014). A concentrated stock solution of PFOA (1×10^−2^ M) was prepared in NERL water and then diluted in NERL water containing 0.5% sucrose. Sucrose is added to the drinking water to ensure consumption of PFOA, but the concentration is significantly lower than the sucrose concentration in sugar-sweetened drinks (10-12%)(Sundborn et al., 2019). This sucrose concentration results in an approximate daily sucrose intake in a 20 g mouse of 21 mg/day, compared to an approximate daily sucrose intake from food of 517 mg/day. Mice were administered vehicle (0.5% sucrose) drinking water or PFOA (8 μM) drinking water *ad libitum* for 6-7 weeks. Food and water consumption were determined on a per cage basis each week and previously reported (Schlezinger et al., 2020). Body weight was measured weekly. Prior to euthanasia, mice were fasted for 6 hours. Three hours into the fast mice were analyzed for body composition using an EchoMRI700 (EchoMRI LLC, Houston, TX) at the Boston University Metabolic Phenotyping Core. Aliquots of liver for lipidomics and gene expression were flash frozen in liquid nitrogen and stored at -80°C. Aliquots of perigonadal adipose samples for histology and gene expression were fixed in 4% paraformaldehyde or flash frozen in liquid nitrogen and stored at -80°C. Blood was collected via cardiac puncture. Serum was separated, flash frozen in liquid nitrogen and stored at -80°C.

### Liver Triacylglyceride Content

Liver triacylglyceride (TG) was assayed as described (Norris et al., 2003). We prepared a saponified neutralized liver extract and created a standard curve with glycerol standard solution (Sigma G7793). Triolein content was assessed using free glycerol reagent (Sigma F6428). Absorbance was read at 540 nM. Triolein was converted to triacylglyceride and divided by wet liver weight to derive liver lipid content (mg TG/ g liver).

### Lipidomics

All serum and liver samples were randomized before sample preparation and again before the analysis.

For liver, the samples were weighed and phosphate-buffered saline (PBS) was added so that the ratio of tissue to buffer was 1 mg tissue to 1 μL buffer. Then, the samples were homogenized manually. 10 μL of each homogenate was taken for the extraction. 400 μL of ice-cold 75% MeOH in water and 200 μL internal standard (c=7 µg/ml) was added to the samples. Tripalmitin-triheptadecanoylglycerol (TG (17:0/17:0/17:0)), purchased from Larodan AB (Solna, Sweden) was used as the internal standard. The samples were then sonicated using a Bandelin Sonorex Digitec (Berlin, Germany) for 10 minutes. After this, 1 mL methyl-tert-butyl ether (MTBE) was added and the samples were shaken for 1 hour, 250 μL HPLC-grade water was added and the samples were then centrifuged (8000 x g, 15 min). An aliquot of 200 μL was taken and transferred to an LC vial and stored at -80 °C until analysis.

For serum lipid analyses, samples were extracted as follows: 10 µL of 0.9% NaCl and 120 µL of CHCl3: MeOH (2:1, v/v) containing internal standards (c=2.5 µg/mL) were added to 10 µL of each plasma sample. The samples were vortex mixed and incubated on ice for 30 min, after which they were centrifuged (9400 x g, 3 min). 60 µL from the lower layer of each sample was then transferred to a glass vial with an insert, and 60 µL of CHCl3: MeOH (2:1, v/v) was added to each sample. The samples were stored at -80 °C until analysis.

The ultra-high-performance liquid chromatography-quadrupole time-of-flight mass spectrometry (UHPLC-QTOFMS) analyses were done in a similar manner as described earlier (McGlinchey et al., 2020). The UHPLC-QTOFMS system was from Agilent Technologies (Santa Clara, CA, USA), combining a 1290 Infinity system and 6545 quadrupole time of flight mass spectrometer, interfaced with a dual jet stream electrospray (dual ESI) ion source. MassHunter B.06.01 software (Agilent Technologies, Santa Clara, CA, USA) was used for all data acquisition and MZmine 2 was used for data processing.

Lipid identification was based on in-house spectral library with retention times. Lipids were normalized with internal standards and (semi) quantitation was performed using lipid-class specific calibration curves. The relative standard deviation of the internal samples in all samples (raw variation) was between 12.4 to 20.5 %, with an average of 17.3%, thus showing the robustness of the method.

### Gene expression analyses

Total RNA was extracted and genomic DNA was removed using the Direct-zol RNA Miniprep Kit (Zymo Research, Orange, CA). cDNA was synthesized from total RNA using the iScript™Reverse Transcription System (BioRad, Hercules, CA). All qPCR reactions were performed using the PowerUp™SYBR Green Master Mix (Thermo Fisher Scientific, Waltham, MA). The qPCR reactions were performed using a StepOnePlus Real-Time PCR System (Applied Biosystems, Carlsbad, CA): UDG activation (50°C for 2 min), polymerase activation (95°C for 2 min), 40 cycles of denaturation (95°C for 15 sec) and annealing (various temperatures for 15 sec), extension (72°C for 60 sec). The primer sequences and annealing temperatures are provided in **Table S3**. Relative gene expression was determined using the Pfaffl method to account for differential primer efficiencies (Pfaffl, 2001), using the geometric mean of the Cq values for beta-2-microglobulin (*B2m*), GAPDH RNA (*Gapdh*) and 18s RNA (*Rn18s*). The average Cq value from two livers or gonadal adipose tissue from female C57/BL6J mice was used as the reference point. Data are reported as “Relative Expression.”

### Histological analyses

5μm adipose tissue sections were stained with hematoxylin and eosin. Micrographs (20x) were visualized on a Nikon Eclipse TE2000 microscope (Nikon Corporation; Tokyo, Japan). Images were analyzed for adipocyte number and size using Adiposoft (Galarraga et al., 2012).

### Statistical analyses

Data are presented as data points from individual mice or as means α standard error (SE). Mice were considered hPPARα positive if they were either homozygous or heterozygous. Information on outliers is presented in Table S1. For statistical analyses of lipidomics data, including Principal Component Analysis (PCA) and hierarchical clustering heatmaps, the data were first log transformed (generalized logarithm transformation) and autoscaled using the Metaboanalyst software (Pang et al., 2020). For evaluation of the changes in the lipid group level, each lipid was first normalized by the mean value of each lipid and the sum of lipids was calculated. To quantify the strengths of associations between lipid groups and clinical variables, Spearman correlation was calculated pairwise between all variables. To visualize this, the *corrplot* R package was used [Simko TWaV. R package “corrplot”: Visualization of a Correlation Matrix. 2017]. In the gene expression analyses, individual values more than four standard deviations different than the mean were considered outliers and were excluded from the analyses. Overall, 10 values were excluded in 8 of the 14 genes analyzed for liver. Within sex and genotype, statistical significance was determined by unpaired, two-tailed t-test (Prism 6, GraphPad Software Inc., La Jolla, CA). Regression analyses were performed using Microsoft R Open version 3.6.1. To investigate the interactions between treatment, sex and genotype in modifying phenotype and gene expression, we used multiple linear regression modeling (MLR). Each outcome was assessed using a MLR model with predictors including sex and an interaction term for genotype and treatment. Models were also stratified by sex, allowing effect estimates to vary between males and females. Statistical significance was evaluated at an α= 0.05 for all analyses.

## Results

We previous reported analyses of PFOA containing drinking water administered to the hPPARα mice and the effect on body weight and composition (Schlezinger et al., 2020). The concentration of PFOA in vehicle drinking water was 94 ng/L and concentrations of PFOA in treatment drinking water averaged 3509 ± 138 μg/L. Both the vehicle and treatment drinking water were found also to contain perfluorohexanoic acid (34 ng/L and 2320 ng/L, respectively). Based on average daily water consumption (0.21 ml/g mouse/day), the daily dose was approximately 0.7 mg/kg/day, resulting in serum concentrations of 47 ± 8 μg/ml in females and 48 ± 10 μg/ml in males. Daily exposure to PFOA for 6 weeks did not significantly impact weight gain in hPPARα mice of either sex. However, PFOA treatment significantly reduced weight gain in male PPARα null mice. A similar trend was observed in female PPARα null mice, but the effect was not significant. Body composition was not affected by PFOA in either genotype or sex. No differences in water or food consumption were observed (not shown). No immunotoxicity was evident as spleen/body weight and thymus/body weight ratios were not significantly different in PFOA treated mice (**Fig. S1**).

PFOA significantly increased liver to body weight ratios in both sexes and both genotypes (**Fig. 1a)**(Schlezinger et al., 2020). PFOA induced a significantly greater effect on liver to body weight ratios in female, but not male, PPARα null mice than in hPPARα mice (**Table 1**). In this 6 week exposure scenario, PFOA (≈ 50 ug/ml serum) induced a significant increase in liver triacylglyceride (TG) in female hPPARα mice (**Fig. 1b**); liver to body weight ratios were highly correlated with liver TG (Pearson r = 0.7696, p = 0.001). In males, PFOA also induced a significant increase in liver TG in hPPARα mice (**Fig. 1b**); liver to body weight ratios were highly correlated with liver TG (Pearson r = 0.8012, p < 0.0001). In females and males, liver TG concentrations were significantly greater in PPARα null mice than in hPPARα mice (**Fig. 1b, Table 1**). PFOA treatment did not have a significant effect on liver TG in female PPARα null mice, and there was a significant interaction between genotype and treatment (**Fig. 1b, Table 1**). There was no significant correlation between liver to body weight ratio and liver TG in female PPARα null mice. PFOA treatment also did not have a significant effect on liver TG in male PPARα null mice (**Fig. 1b**), but there was a trend toward a correlation between liver to body weight ratios and liver TG (Pearson r = 0.5439, p =0.08).

**Table 1:** Effect estimates (β) and standard errors (SE) for liver characteristics. Regression models were fit to evaluate associations of phenotypic outcomes with treatment and genotype, including a treatment-genotype interaction term. The left hand column adjusts for sex. The two right columns stratify by sex, allowing results to differ between males and females. Statistical significance was evaluated at α = 0.05 for all analyses. Liver/body weight data were previously published (Schlezinger et al., 2020).

| Test | ALL |  | MALE |  | FEMALE |  |
| --- | --- | --- | --- | --- | --- | --- |
| | $\beta$ (SE) | P value | $\beta$ (SE) | P value | $\beta$ (SE) | P value |
| **Liver to Body Weight (%)** |  |  |  |  |  |  |
| *PFOA treatment* | 4.75 (0.39) | **<0.0001** | 4.99 (0.63) | **<0.0001** | 4.49 (0.44) | **<0.0001** |
| *hPPARα Genotype* | -0.41 (0.36) | 0.25 | -0.19 (0.54) | 0.72 | -0.60 (0.43) | 0.17 |
| *Treatment\*Genotype* | -1.17 (0.51) | **0.03** | -0.95 (0.79) | 0.24 | -1.50 (0.60) | **0.02** |
| *Male Sex* | 0.43 (0.25) | 0.10 | - | - | - | - |
| **Liver Triacylglycerides (mg/g liver)** |  |  |  |  |  |  |
| *PFOA treatment* | 6.18 (4.42) | 0.17 | 15.04 (7.68) | 0.06 | -1.93 (3.08) | 0.54 |
| *hPPARα Genotype* | -22.04 (4.04) | **<0.0001** | -21.14 (6.68) | **0.004** | -21.96 (2.97) | **<0.0001** |
| *Treatment\*Genotype* | 9.96 (5.79) | 0.09 | 8.30 (9.73) | 0.40 | 8.80 (4.20) | **0.048** |
| *Male Sex* | 5.44 (2.87) | 0.06 | - | - | - | - |

**Figure 1.**
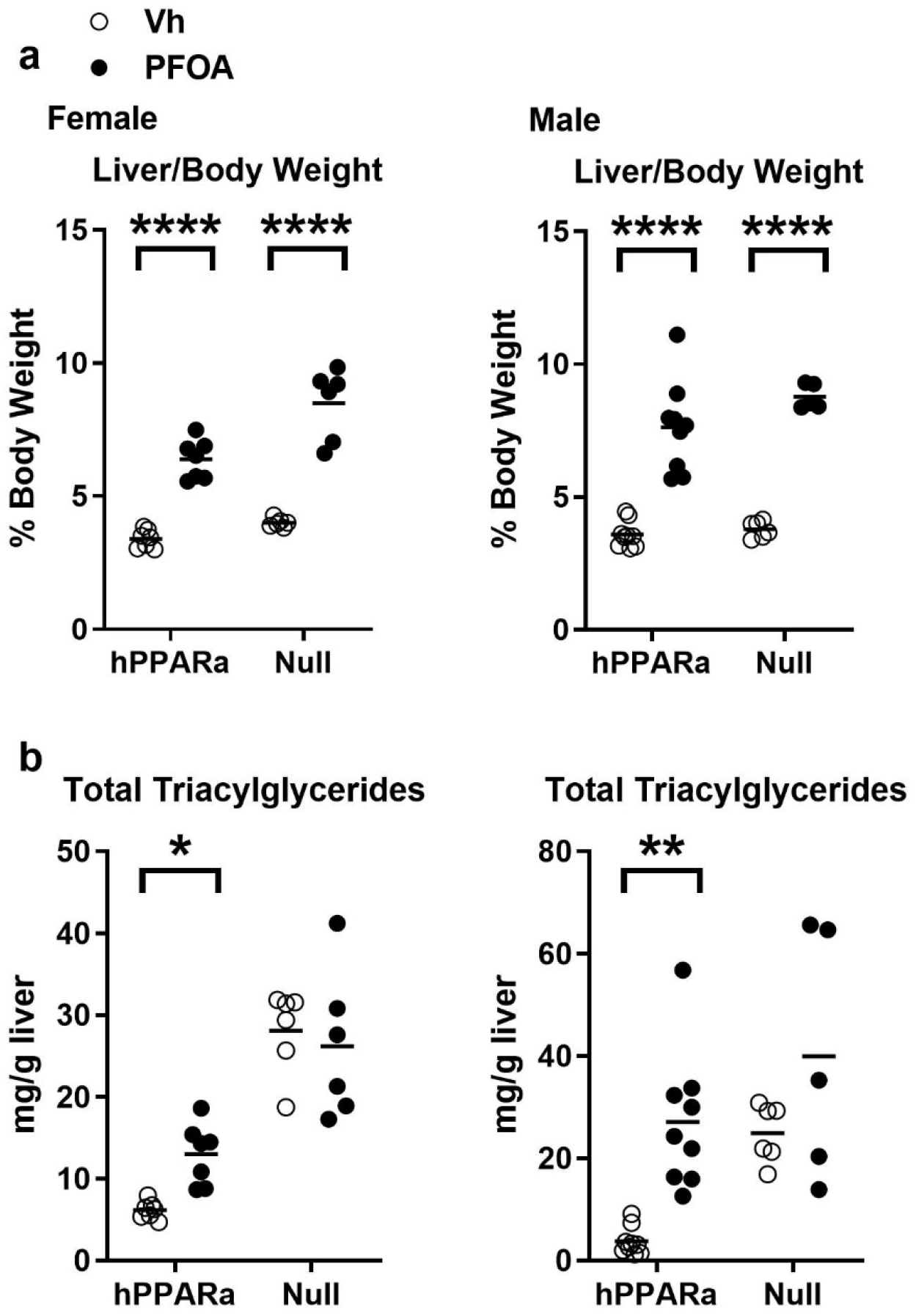
Liver weight and triacylglyceride concentrations in PFOA-exposed mice. Three-week-old male and female hPPARα and PPARα null mice were treated with either vehicle (Vh, NERL water with 0.5% sucrose) or PFOA (8 µM in NERL water with 0.5% sucrose) in drinking water for 6 weeks. During treatment, the mice were fed an American Diet (see Table S1). **a** Liver to body weight ratios were determined at euthanasia (previously reported in Schlezinger et al., 2020). **b** Total triacylglyceride concentrations were determined from measuring triolein content in saponified neutralized liver extracts. Data are from individual mice, with the mean indicated by a line. N = 5-9. Significantly different from Vh (* p<0.05, ** p<0.01, **** p<0.0001, t-test).

Genotype and PFOA treatment had a significant effect on the composition of TG in the liver. The PCA analysis shows, regardless of sex, that lipid profiles cluster for livers from Vh and PFOA mice in a genotype-distinct manner (**Fig. 2a**). PPARα genotype had a major impact on the lipidome, with over 30% of the identified lipids showing significant differences (p, FDR <0.05)(**Fig. 2b, Fig. S2, Tables S4-5**). There were significantly lower levels of the majority of TG in the hPPARα mice compared to PPARα null mice (**Fig. 3, Table S4**). PFOA exposure had a very significant impact on the liver lipidome and a significant interaction with genotype, with more a drastic impact in the hPPARα mice (24% of lipids), and of those, the impact was more drastic in male mice (14% vs 0.4%, significantly different lipids, male vs female, respectively)(**Fig. 2b, Table S4**). TG were increased by PFOA exposure in hPPARα mice (**Fig. 3, Table S4**). The increases were largely attributable to increases in TG substituted with saturated and monounsaturated fatty acids, but not polyunsaturated fatty acids, with same trends in both sexes (**Fig. 3, Table S4-5**). The intra-TG class correlations in hPPARα mice were strengthened by PFOA treatment overall (**Fig. S3a-b**). The effect of PFOA on TG was mitigated in PPARα null mice. Interestingly, the impact of the PFOA exposure in PPARα null mice, being clearly less drastic (7% of lipids) overall, was much more drastic in female mice (8.5% of lipids) than males, although the patterns of effect were the same (**Fig. 3, Table S4-5**). The intra-TG class correlation in PPARα null mice was showing an opposite trend, with PFOA treatment strengthening the correlations (**Fig. S3c-d**). Overall, genotype and PFOA treatment were more important predictors of TG composition than sex.

**Figure 2.**
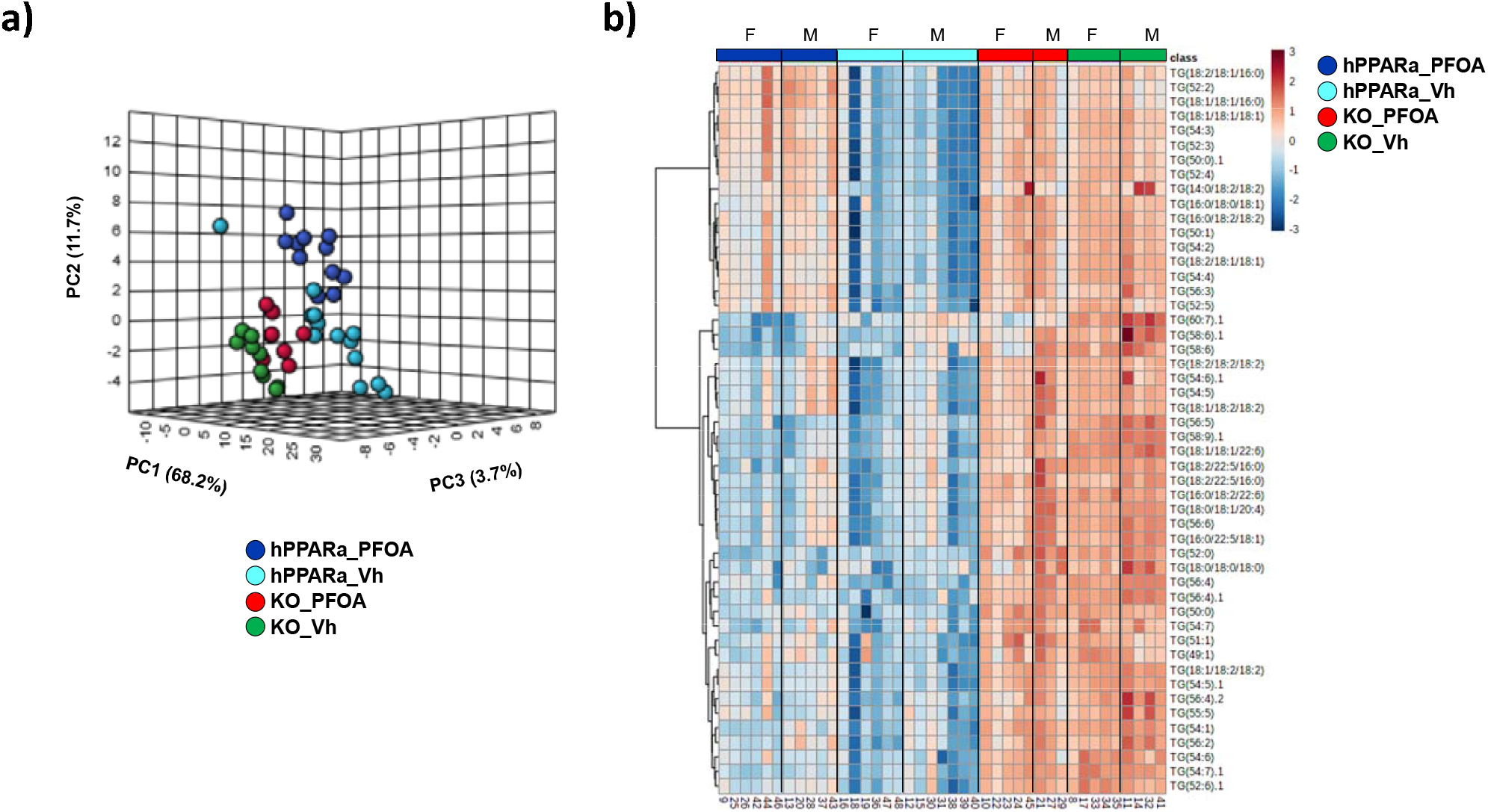
Triacylglyceride profiles of liver. hPPARα and PPARα null mice were treated with vehicle or PFOA in drinking water for 6 weeks, as described in Fig. 1. TGs were identified and quantified by UHPLC-QTOFMS. **a** Principal component analysis. **b** Heatmap of top 50 TGs selected by ANOVA analysis. N = 3-6.

**Figure 3.**
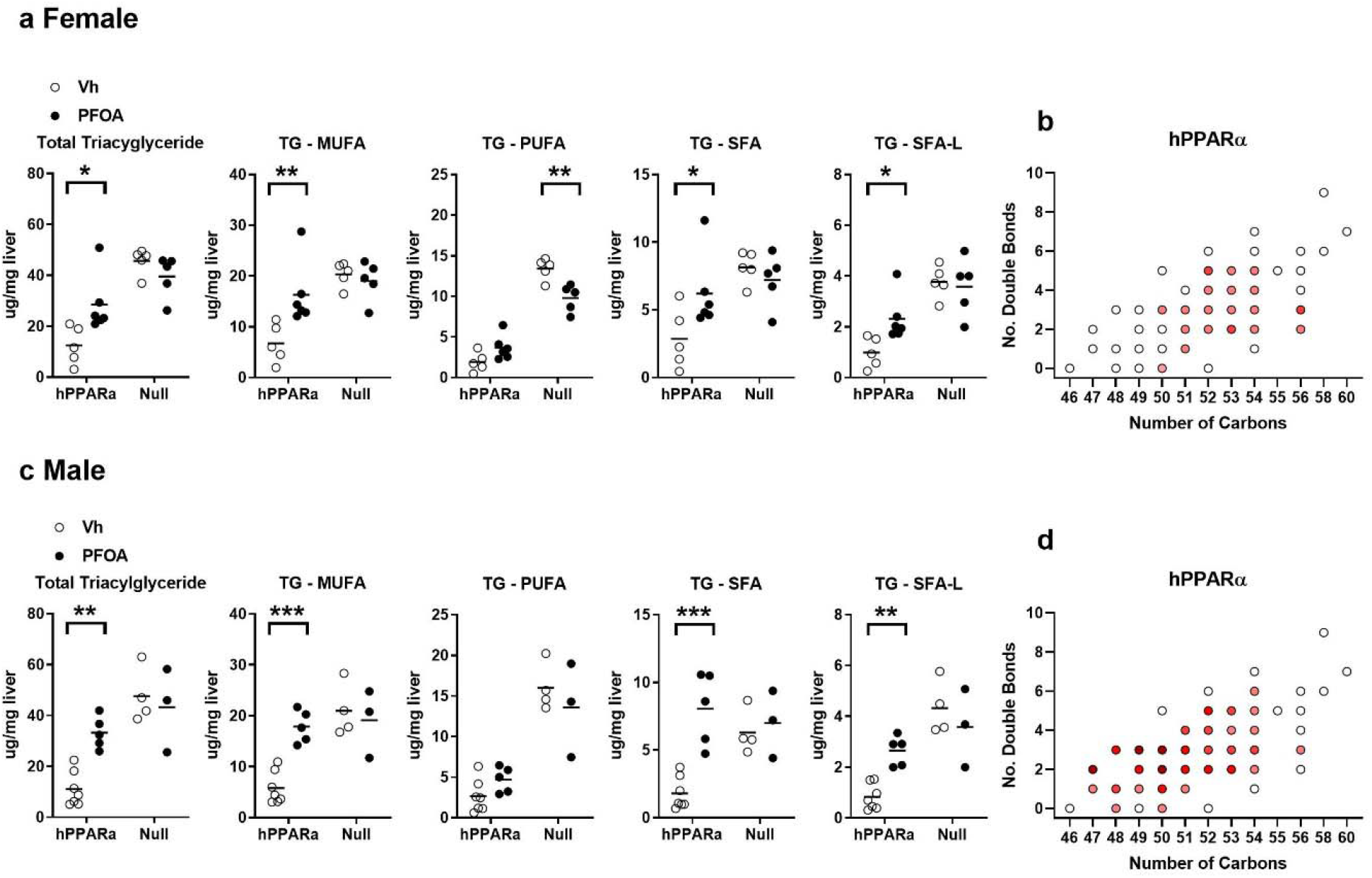
Quantification of triacylglyceride types in liver. hPPARα and PPARα null mice were treated with vehicle or PFOA in drinking water for 6 weeks, as described in Fig. 1. TGs were identified and quantified by UHPLC-QTOFMS. **a** Analysis of select lipids in females. **b** Analysis of select lipids in males. Data are from individual mice, with the mean indicated by a line. N = 3-6. Significantly different from Vh (* p<0.05, ** p<0.01, *** p<0.001, **** p<0.0001, t-test).

We previously showed that PFOA significantly increased cholesterol associated with lipoprotein particles, particularly in male mice, with a shift in the distribution from HDL-c to very low-density lipoprotein-c (vLDL-c) and LDL-c (Schlezinger et al., 2020). Here, we investigated the concentration and distribution of TG within lipoprotein particles (**Fig. 4**). In females, PFOA increased the amount of TG associated with lipoproteins in hPPARα mice by 1.8-fold, which was associated with an increase in the proportion of vLDL-c (**Fig. 4a, 4b**). The amount of TG associated with lipoproteins in untreated female PPARα null mice was 2.2-fold higher than that in untreated hPPARα mice and was increased further (1.6-fold) in PFOA-treated PPARα null mice (**Fig. 4a, 4b**). In males, PFOA modestly decreased the amount of TG associated with lipoproteins in hPPARα mice by 0.3-fold; however, the proportion associated with vLDL-c increased (**Fig. 4c, 4d**). The amount of TG associated with lipoproteins in untreated male PPARα null mice was 1.7-fold higher than that in untreated hPPARα mice, and this was modestly decreased (0.2 fold) in PFOA-treated PPARα null mice (**Fig. 4c, 4d**). The distribution of TG across the lipoprotein particle types was unchanged in PFOA treated male PPARα null mice (**Fig. 4c, 4d**).

**Figure 4.**
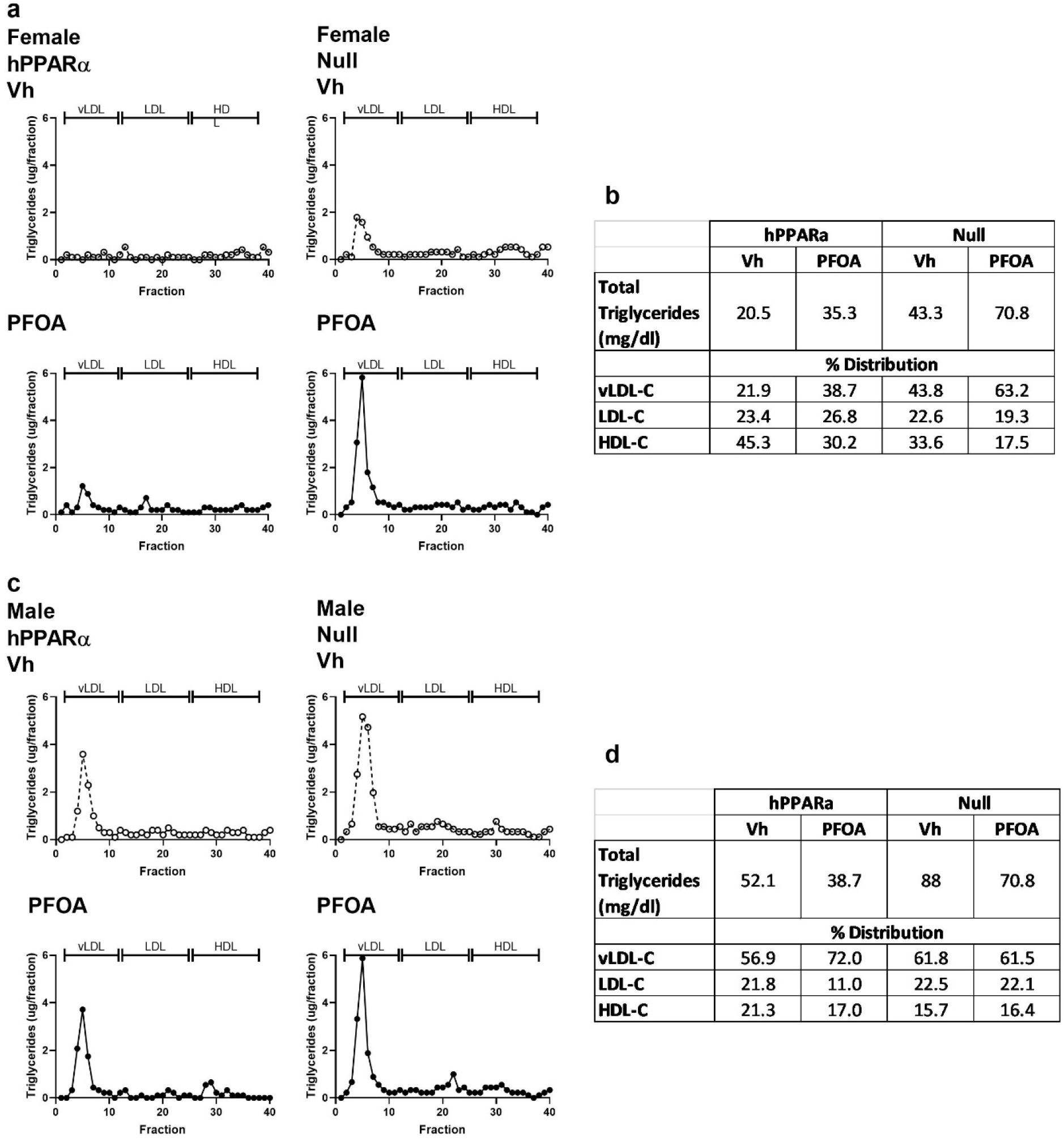
Amount and distribution of triacylglycerides associated with lipoprotein particles in PFOA-exposed mice. hPPARα and PPARα null mice were treated with vehicle or PFOA in drinking water for 6 weeks, as described in Fig. 1. Serum was pooled into a single sample per group, and lipoproteins were separated by FPLC. Triacylglycerides in each fraction were determined enzymatically. **a** Female profile. **b** Quantification of in triacylglycerides and distribution across lipoprotein particles in females. **c** Males profile. **d** Quantification of total triacylglycerides and distribution across lipoprotein particles in males.

Genotype, sex, and PFOA treatment all had a significant effect on the composition of serum TGs. The PCA analysis showed that lipid profiles clustered for serum from Vh and PFOA mice in a genotype- and sex-distinct manner (**Fig. 5a**). Interestingly, sex had a very strong impact on the serum lipidome, while genotype had much less impact (60% vs 13% of lipids significantly different, sex vs genotype)(**Fig. 5b, Tables S6-7**). There were significantly lower levels of the majority of TG in female hPPARα mice compared to males PPARα mice, but this was blunted in PPARα null mice (**Fig. 3, Table S4**). PFOA exposure also had a significant impact on serum lipidome (**Fig. 5b, Table S6**). In this 6 week exposure scenario, PFOA (≈ 50 ug/ml serum) induced little change in total serum TG concentrations in hPPARα mice of either sex (**Fig. 6, Table S6**). Interestingly, distinctly from the liver, PFOA had a stronger effect in PPARα null mice than hPPARα mice. In female PPARα null mice (**Fig. 6a, Tables S6-7**), PFOA significantly increased serum TG, with significant increases in TG substituted with saturated and monounsaturated fatty acids, but not polyunsaturated fatty acids. A similar trend was observed in PPARα null males, but the increases did not reach significance (**Fig. 6b, Tables S6-7**). Overall, genotype, sex and PFOA treatment were all important predictors of TG composition.

**Figure 5.**
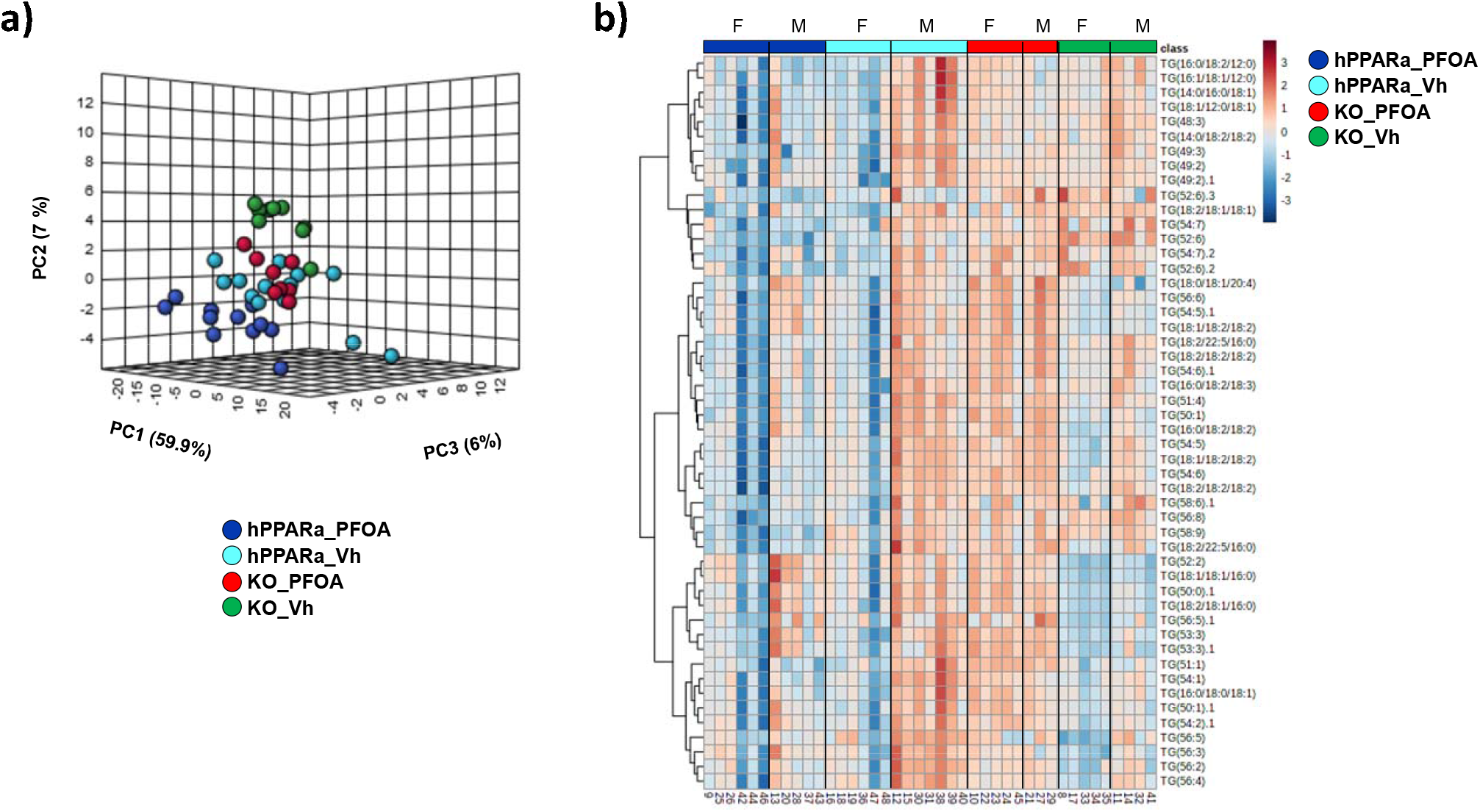
Triacylglyceride profiles of serum. hPPARα and PPARα null mice were treated with vehicle or PFOA in drinking water for 6 weeks, as described in Fig. 1. TGs were identified and quantified by UHPLC-QTOFMS. **a** Principal component analysis. **b** Heatmap of top 50 TGs selected by ANOVA analysis N = 3-6.

**Figure 6.**
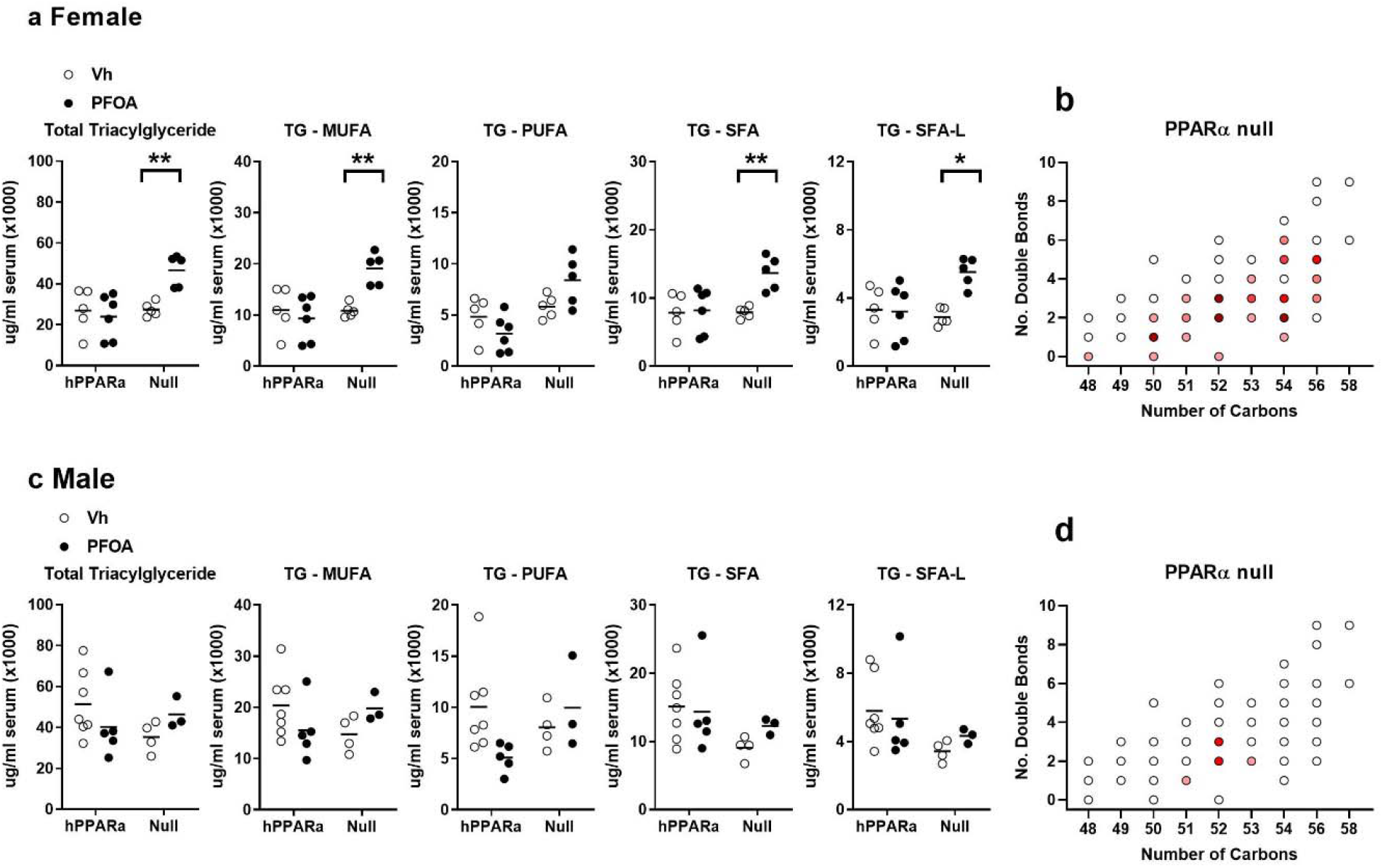
Quantification of triacylglyceride types in serum. hPPARα and PPARα null mice were treated with vehicle or PFOA in drinking water for 6 weeks, as described in Fig. 1. Lipids were identified and quantified by UHPLC-QTOFMS. **a** Analysis of select lipids in females. **b** Analysis of select lipids in males. Data are from individual mice, with the mean indicated by a line. N = 3-6. Significantly different from Vh (* p<0.05, ** p<0.01, *** p<0.001, **** p<0.0001, t-test).

To begin to investigate the pathways contributing to the changes in TG homeostasis we observed in the liver and serum, we quantified the expression of genes of transcriptional regulators of TG homeostasis (**Fig. 7**). PGC1α and lipin-1 coactivate PPARα and play important roles in enhancing mitochondrial fatty acid oxidation (Finck et al., 2006; Vega et al., 2000). Loss of lipin-1 increases circulating TG levels (Chen et al., 2008; Finck et al., 2006). In female mice, PFOA did not change *Pgc1a* expression in hPPARα mice but significantly decreased *Pgc1a* expression in PPARα null mice and with a significant interaction between treatment and genotype (**Fig. 7a, Table 2**). PFOA significantly decreased expression of *Lpin1* in both hPPARα and PPARα null female mice (**Fig. 7a, Table 2**). In contrast, PFOA had no effect on expression of *Pgc1a* and *Lpin1* in either hPPARα or PPARα null male mice (**Fig. 7b, Table 2**).

**Table 2:**
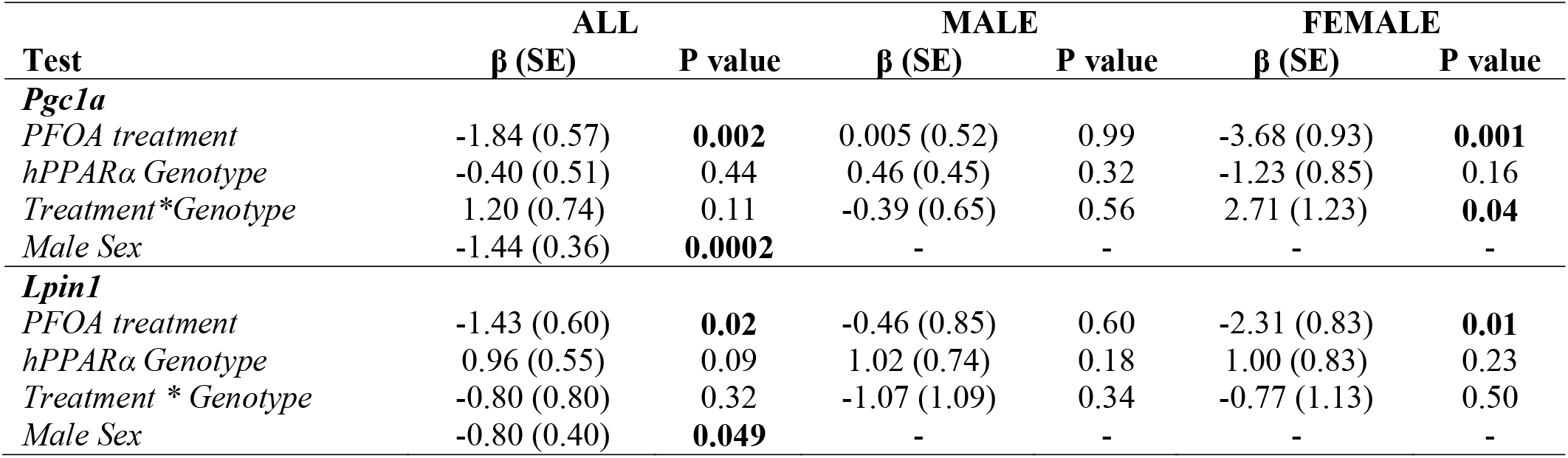
Effect estimates (β) and standard errors (SE) for liver expression of transcriptional modulators. Regression models were fit to evaluate associations of gene expression outcomes with treatment and genotype, including a treatment-genotype interaction term. The left-hand column adjusts for sex. The two right columns stratify by sex, allowing results to differ between males and females. Statistical significance was evaluated at α = 0.05 for all analyses.

**Figure 7.**
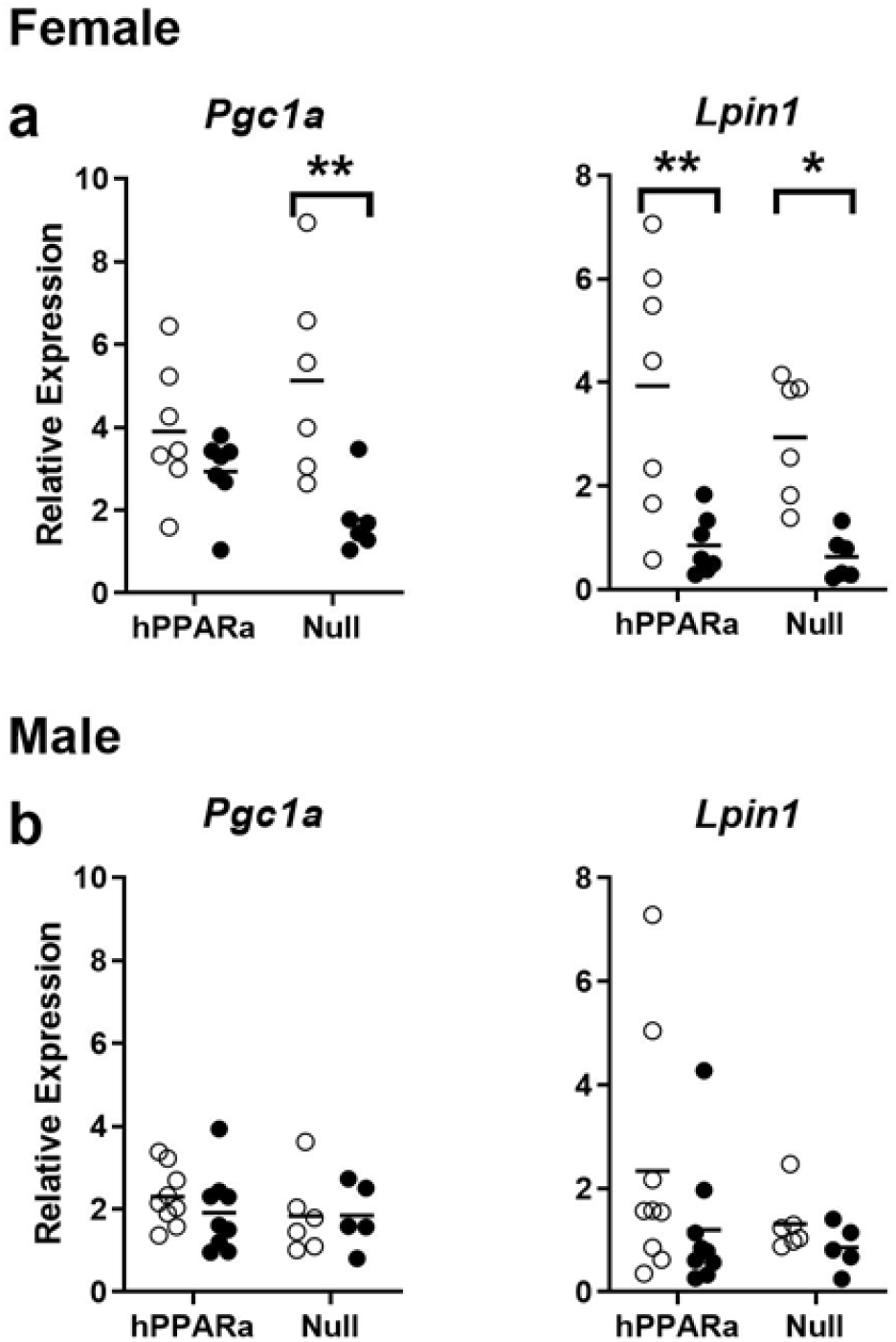
Expression of transcriptional regulators in liver of PFOA-exposed mice. hPPARα and PPARα null mice were treated with vehicle or PFOA in drinking water for 6 weeks, as described in Fig. 1. Following isolation of RNA from liver, gene expression was determined by RT-qPCR. **a** Female mice. **b** Male mice. Data are from individual mice, with the mean indicated by a line. N = 5-9. Significantly different from Vh (* p<0.05, ** p<0.01, *** p<0.001, **** p<0.0001, t-test).

Triacylglyceride concentrations in liver and serum are determined by multiple processes (**Fig. 8**): uptake of fatty acids, fatty acid synthesis, assembly and secretion of cholesterol particles, lipolysis and fatty acid oxidation (Alves-Bezerra and Cohen, 2017). We investigated the effect of PFOA on the expression of biomarker genes for each of these processes in liver. CD36 is a receptor that carries out uptake of fatty acids into the liver. PFOA significantly increased expression of *Cd36* in female and male hPPARα mice, and this increase was strongly reduced in PPARα null mice (**Fig. 9a, Table 3**). Fatty acid synthase is a rate limiting enzyme in *de novo* fatty acid synthesis. PFOA did not affect *Fasn* gene expression in either hPPARα or PPARα null female mice (**Fig. 9b, Table 3**). However, PFOA significantly increased *Fasn* expression in male hPPARα mice but not in male PPARα null mice (**Fig. 9b, Table 3**). Acyl-CoA derivatives of fatty acids are substrates for both lipid synthesis and oxidation, and long-chain acyl-CoA synthetases catalyze the formation of acyl-CoAs. PFOA significantly increased expression of *Acsl5* in female and male hPPARα mice. The increase was only significantly abrogated in female PPARα null mice (**Fig. 9c, Table 3**). TG are formed by the esterification of fatty acyl-CoAs to glycerol. MOGAT1 catalyzes the synthesis of diacylglycerols, and DGAT1 catalyzes the acylation of diacylglcerol. PFOA significantly increased expression of *Mogat1* in female and male hPPARα mice, with a significantly greater induction occurring in male than in female mice (**Fig. 9d, Table 3**). Induction of *Mogat1* by PFOA was strongly reduced in PPARα null mice (**Fig. 9d, Table 3**). In contrast, *Dgat1* was only modestly induced by PFOA in female hPPARα mice (**Fig. 9d, Table 3**). PFOA had no effect on *Dgat1* expression in female PPARα null, male PPARα null or male hPPARα mice (**Fig. 9d, Table 3**).

**Table 3:**
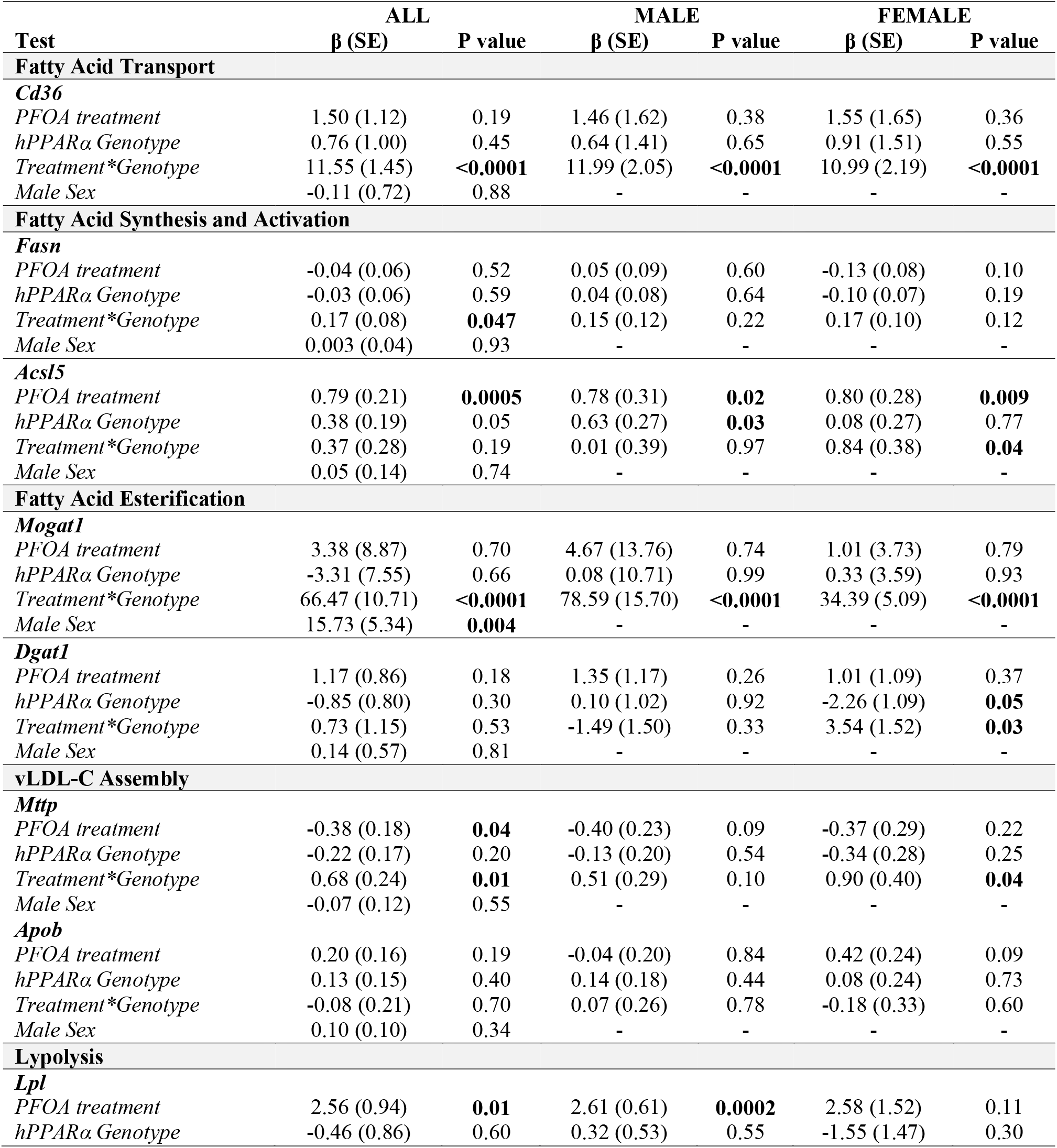
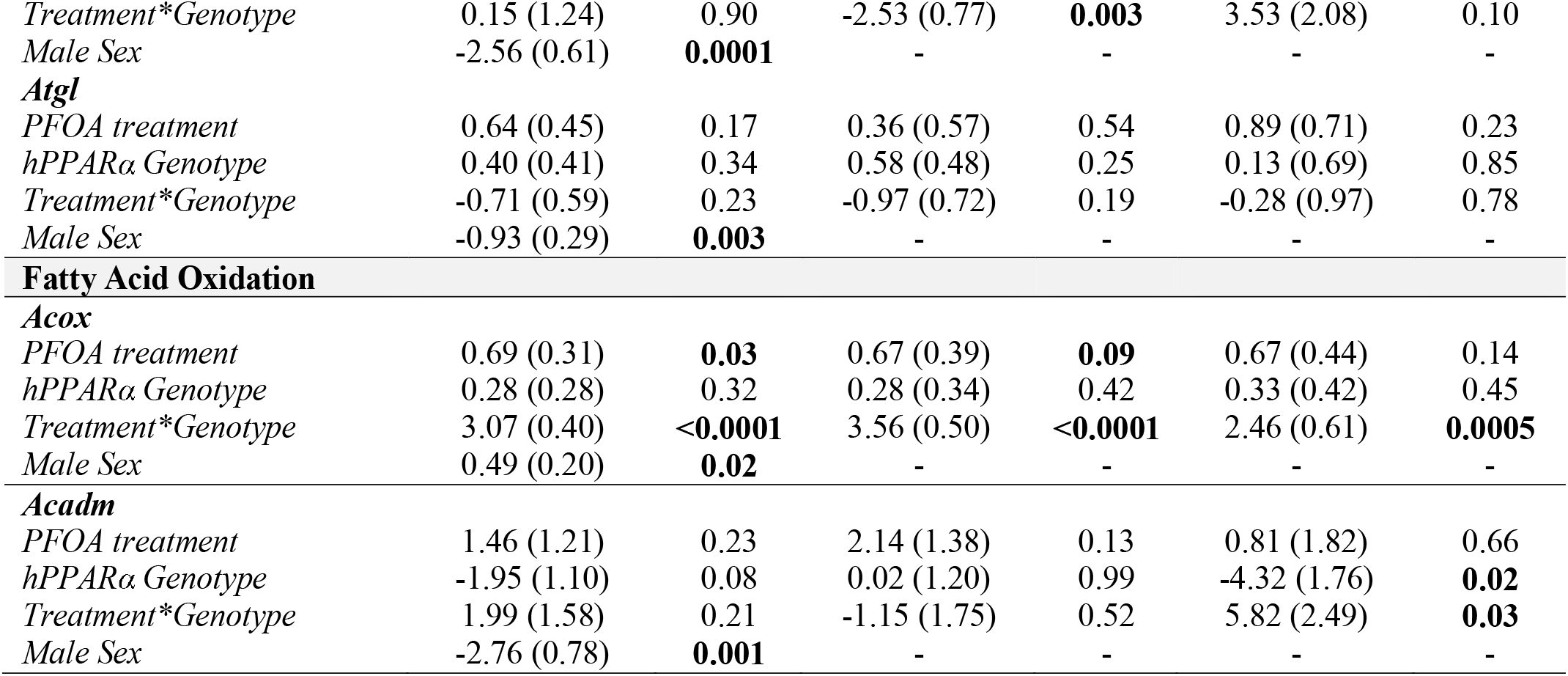
Effect estimates (β) and standard errors (SE) for liver expression of gene involved in regulating lipid homeostasis. Regression models were fit to evaluate associations of gene expression outcomes with treatment and genotype, including a treatment-genotype interaction term. The left-hand column adjusts for sex. The two right columns stratify by sex, allowing results to differ between males and females. Data for *Cd36, Mogat1, Apob* and *Acox* were previously reported (Schlezinger et al., 2020) Statistical significance was evaluated at α = 0.05 for all analyses.

**Figure 8.**
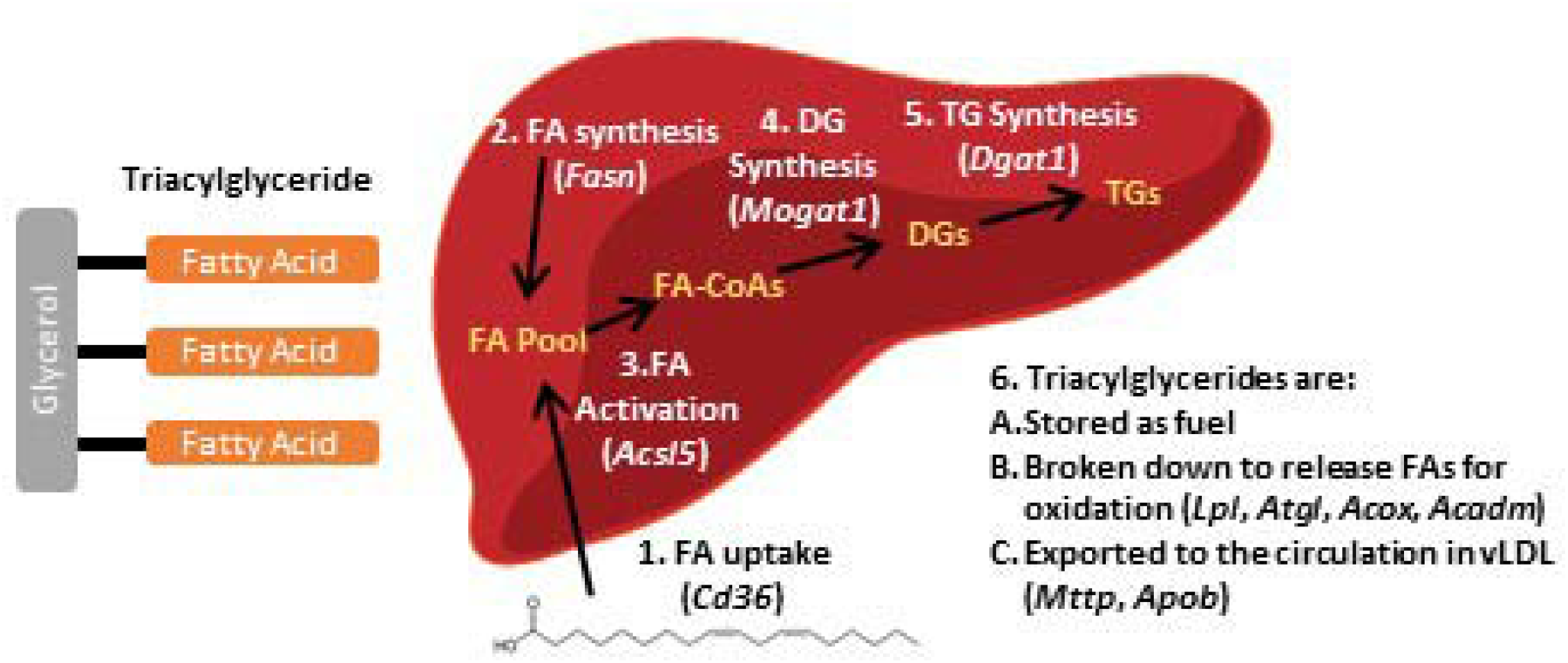
Triacylglyceride synthesis and use pathways in liver. FA: fatty acid, FA-CoA: fatty acid-CoA, DG: diacylglyceride, TG: triacylglyceride.

**Figure 9.**
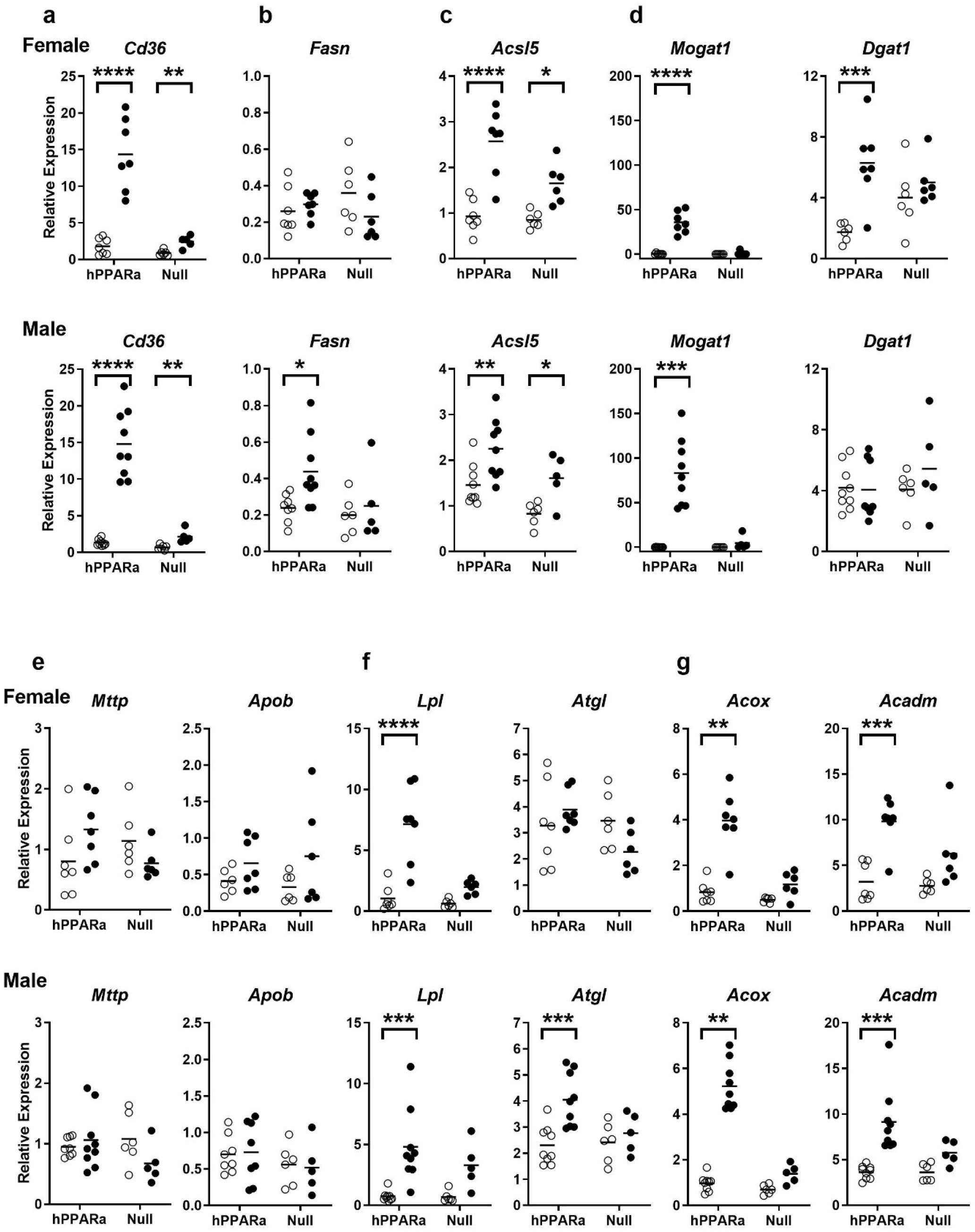
Fatty acid and triacylglyceride homeostasis-related gene expression in liver of PFOA-exposed mice. hPPARα and PPARα null mice were exposed to Vh or PFOA in their drinking water for 6 weeks, as described in Fig. 1. Following isolation of RNA from liver, gene expression was determined by RT-qPCR. Expression of biomarker genes for the following processes were quantified: **a** Fatty acid uptake. **b** Fatty acid synthesis. **c** Fatty acid activation. **d** Fatty acid esterification. **e** Triacylglyceride export. **f** Lipolysis. **g** Fatty acid oxidation. *Acox, Apob, CD36* and *Mogat1* data were previously reported (Schlezinger et al., 2020). Data are from individual mice, with the mean indicated by a line. N = 5-9. Significantly different from Vh (* p<0.05, ** p<0.01, *** p<0.001, **** p<0.0001, t-test).

Assembly and secretion of vLDL requires TG to be incorporated into the lipoprotein ApoB100 by microsomal TG transfer protein as ApoB100 is being translated in the ribosome. While PFOA did not induce a significant change *Mttp* expression in either sex individually, (**Fig. 9e**), when considered together, PFOA induced a hPPARα-dependent increase in *Mttp* expression (**Table 3**). PFOA induced no significant changes in either *Apob* expression in either sex or genotype (**Fig. 9e, Table 3**).

Lipolysis, carried out by enzymes in the liver including lipid droplet-associated adipose triacylglyceride lipase and in the serum, including lipoprotein lipase, release fatty acids from TG molecules. PFOA significantly increased expression of *Lpl* in female and male hPPARα mice, with a significantly greater increase in female than in male mice (**Fig. 9f, Table 3**). These increases were strongly reduced in PPARα null mice (**Fig. 9f, Table 3**). In contrast, *Atgl* was only induced by PFOA in male and not female hPPARα mice (**Fig. 9f, Table 3**). *Atgl* was expressed to a lesser extent in male than in female mice (**Table 3**).

Finally, fatty acid oxidation is catalyzed by peroxisomal and mitochondrial enzymes such as acyl-CoA oxidase and acyl-coA dehydrogenase medium chain, respectively. PFOA significantly increased expression of *Acox* and *Acadm* in female and male hPPARα mice, and these increases were strongly reduced in PPARα null mice (**Fig. 9g, Table 3**). Overall, male mice expressed a higher level of *Acox*, while female mice expressed a higher level of *Acadm* (**Table 3**).

*Mogat1* is a PPARα regulated gene (Lutkewitte et al., 2019). In this study, *Mogat1* expression was highly correlated in females with expression of *Cd36, Acsl5, Lpl, Acox*, and *Acadm* (Pearson r > 0.7 with p ≤ 0.0001 females). Female *Mogat1* expression was not correlated with *Pgc1a, Lpin1, Fasn, Dgat1, Mttp, Apob*, or *Atgl*. In males, *Mogat1* expression was highly correlated with expression of *Cd36, Acox*, and *Acadm* (Pearson r > 0.58 with p ≤ 0.001) and modestly correlated with expression of *Atgl, Fasn, Acsl5*, and *Apob* (Pearson r = 0.4-0.5 with p ≤ 0.03). Male *Mogat1* expression was not correlated with expression of *Pgc1a, Lpin1, Dgat1, Mttp*, or *Lpl*.

Adipose tissue is a significant source of fatty acids, and thus, contributor to triacylglyceride homeostasis in liver (Alves-Bezerra and Cohen, 2017). We previously reported that PFOA did not have a significant effect on adipose amount in either sex or in either hPPARα and PPARα null mice (Schlezinger et al., 2020). Here, we examined the effect of PFOA on adipose histology and gene expression using the perigonadal fat pad, because we hypothesized that PFOA could alter the supply of fatty acids from the adipose to the liver. PFOA did not have a significant effect on the relative weight of this fat pad, the number of adipocytes or the size of the adipocytes in either sex or genotype (**Fig. 10a-d**). Overall, males had a larger perigonadal fat pad than females (**Fig. 10b, 10d, 10e**). In addition, the weight of this fat pad was highly correlated with total weight gain in males (hPPARα: Pearson r = 0.7189, p = 0.0008; Null: Pearson r = 0.8212, p = 0.0019). In females, fat pad weight was correlated with total weight gain in null but not hPPARa mice (hPPARα: Pearson r = 0.3075, p = 0.25; Null: Pearson r = 0.7048, p = 0.01).

**Figure 10.**
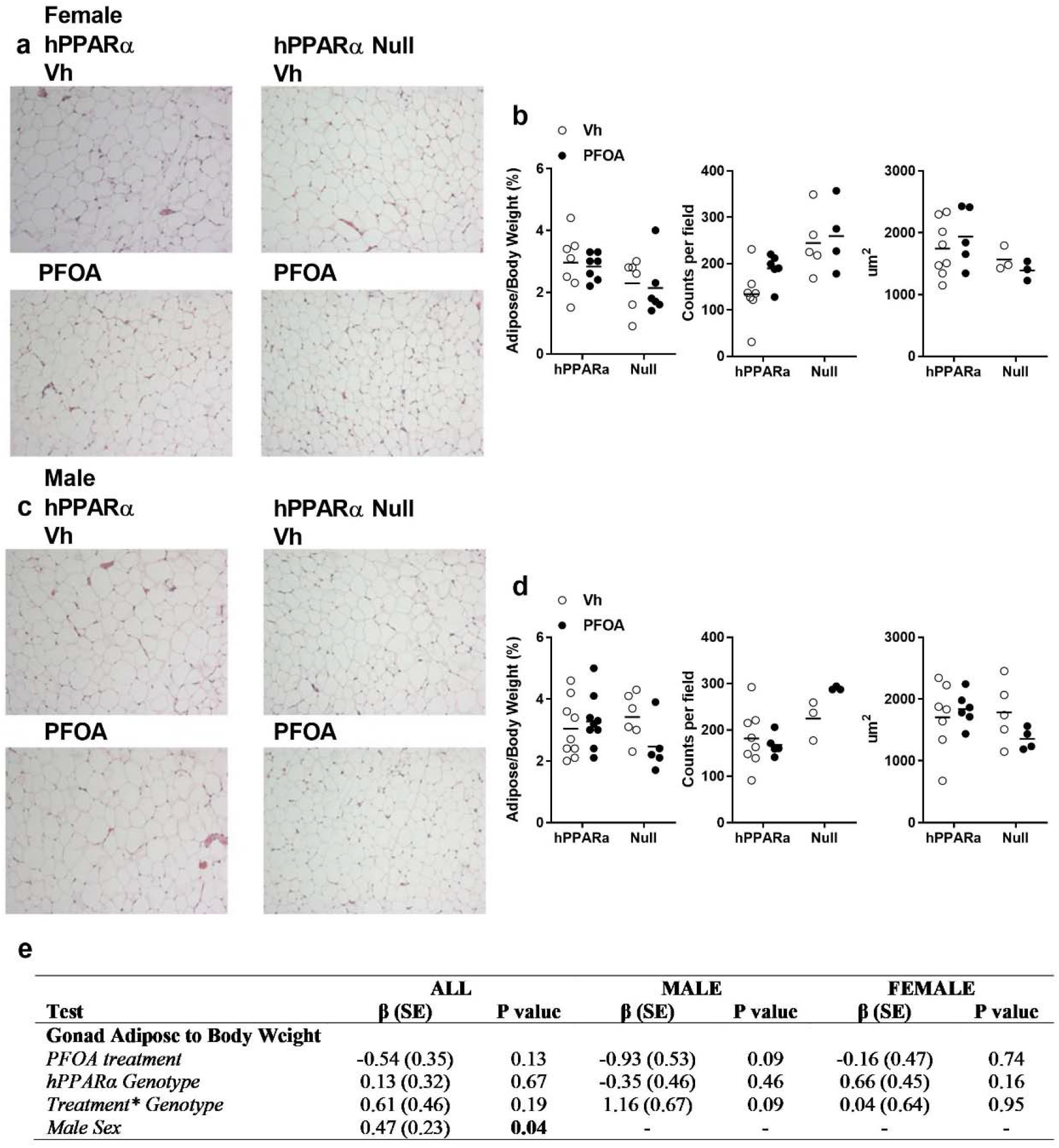
PFOA does not change adiposity in hPPARα or PPARα null mice. hPPARα and PPARα null mice were exposed to Vh or PFOA in their drinking water for 6 weeks, as described in Fig. 1. **a, c** H&E staining of perigonadal adipose sections. **b, d** Analyses of perigonadal adipose. **e**. Results of multiple linear regression analysis of relative adipose weight. Data are from individual mice, with the mean indicated by a line. N = 3-9.

Next, we investigated whether PFOA induced changes in adipocyte gene expression. As noted above, PGC1α and lipin-1 play important roles in regulating fatty acid and TG homeostasis in liver (Puigserver et al., 1998), and this is also true in adipose (Nadra et al., 2012). Fatty acid oxidation is an essential feature of brown/brite adipocytes (Calderon-Dominguez et al., 2016), and *Acox* is a well-known transcriptional target of PPARα. PNPLA2 (encoded by *Atgl*) liberates fatty acids from TG. Adiponectin and leptin are adipokines that play an important role in metabolic homeostasis, and a decreasing adiponectin/leptin ratio is used as a biomarker of adipose dysfunction in humans (Frühbeck et al., 2018). In our hands, PFOA did not induce changes in expression of *Pgc1a, Lpin1, Acox, Atgl, Adipoq* or *Lep* (**Fig. 11, Table 4**).

**Table 4:** Effect estimates (β) and standard errors (SE) for adipose expression of gene involved in regulating lipid homeostasis. Regression models were fit to evaluate associations of gene expression outcomes with treatment and genotype, including a treatment-genotype interaction term. The left-hand column adjusts for sex. The two right columns stratify by sex, allowing results to differ between males and females. Statistical significance was evaluated at α = 0.05 for all analyses.

| Test | ALL |  | MALE |  | FEMALE |  |
| --- | --- | --- | --- | --- | --- | --- |
| | $\beta$ (SE) | P value | $\beta$ (SE) | P value | $\beta$ (SE) | P value |
| ***Pgc1a*** |  |  |  |  |  |  |
| *PFOA treatment* | -0.06 (0.10) | 0.57 | -0.15 (0.13) | 0.26 | 0.07 (0.15) | 0.65 |
| *hPPARα Genotype* | -0.07 (0.10) | 0.51 | 0.06 (0.15) | 0.67 | -0.18 (0.15) | 0.24 |
| *Treatment\*Genotype* | -0.02 (0.15) | 0.88 | -0.08 (0.22) | 0.71 | -0.02 (0.21) | 0.92 |
| *Male Sex* | 0.04 (0.08) | 0.62 | - | - | - | - |
| ***Lpin1*** |  |  |  |  |  |  |
| *PFOA treatment* | 0.27 (0.34) | 0.44 | 0.04 (0.47) | 0.93 | 0.60 (0.49) | 0.23 |
| *hPPARα Genotype* | 0.18 (0.37) | 0.63 | -0.42 (0.54) | 0.44 | 0.78 (0.48) | 0.13 |
| *Treatment\*Genotype* | -0.22 (0.54) | 0.68 | 0.84 (0.79) | 0.31 | -1.23 (0.71) | 0.09 |
| *Male Sex* | 0.15 (0.27) | 0.57 | - | - | - | - |
| ***Plin1*** |  |  |  |  |  |  |
| *PFOA treatment* | 0.27 (0.34) | 0.44 | 0.04 (0.47) | 0.94 | 0.60 (0.48) | 0.23 |
| *hPPARα Genotype* | 0.18 (0.37) | 0.63 | -0.42 (0.54) | 0.44 | 0.78 (0.48) | 0.13 |
| *Treatment\*Genotype* | -0.22 (0.54) | 0.68 | 0.84 (0.80) | 0.31 | -1.23 (0.71) | 0.09 |
| *Male Sex* | 0.15 (0.27) | 0.76 | - | - | - | - |
| ***Acox*** |  |  |  |  |  |  |
| *PFOA treatment* | 0.26 (0.70) | 0.71 | -0.58 (1.06) | 0.59 | 1.48 (0.86) | 0.10 |
| *hPPARα Genotype* | 0.06 (0.76) | 0.94 | -0.92 (1.21) | 0.46 | 1.18 (0.86) | 0.19 |
| *Treatment\*Genotype* | -1.00 (1.11) | 0.37 | 0.87 (1.80) | 0.64 | -3.05 (1.24) | **0.02** |
| *Male Sex* | 1.49 (0.56) | **0.01** | - | - | - | - |
| ***Atgl*** |  |  |  |  |  |  |
| *PFOA treatment* | 0.19 (0.46) | 0.68 | -0.16 (0.56) | 0.78 | 0.70 (0.72) | 0.34 |
| *hPPARα Genotype* | 0.58 (0.49) | 0.25 | -0.41 (0.64) | 0.53 | 1.56 (0.72) | **0.04** |
| *Treatment\*Genotype* | -0.34 (0.73) | 0.64 | 1.37 (0.96) | 0.16 | -1.97 (1.05) | 0.08 |
| *Male Sex* | -0.43 (0.36) | 0.24 | - | - | - | - |
| ***Adipoq*** |  |  |  |  |  |  |
| *PFOA treatment* | -0.02 (0.09) | 0.79 | -0.01 (0.11) | 0.93 | -0.05 (0.17) | 0.78 |
| *hPPARα Genotype* | 0.13 (0.10) | 0.19 | 0.25 (0.12) | 0.05 | 0.01 (0.17) | 0.93 |
| *Treatment \* Genotype* | -0.14 (0.15) | 0.35 | -0.22 (0.18) | 0.25 | -0.06 (0.25) | 0.81 |
| *Male Sex* | -0.26 (0.07) | **0.001** | - | - | - | - |
| ***Lep*** |  |  |  |  |  |  |
| *PFOA treatment* | -0.36 (0.76) | 0.64 | -1.26 (1.23) | 0.32 | 0.97 (0.67) | 0.16 |
| *hPPARα Genotype* | -0.41 (0.82) | 0.62 | -1.76 (1.41) | 0.22 | 1.24 (0.67) | 0.08 |
| *Treatment\*Genotype* | -0.27 (1.20) | 0.82 | 0.51 (2.09) | 0.81 | -1.52 (0.97) | 0.13 |
| *Male Sex* | 1.35 (0.59) | **0.03** | - | - | - | - |

**Figure 11.**
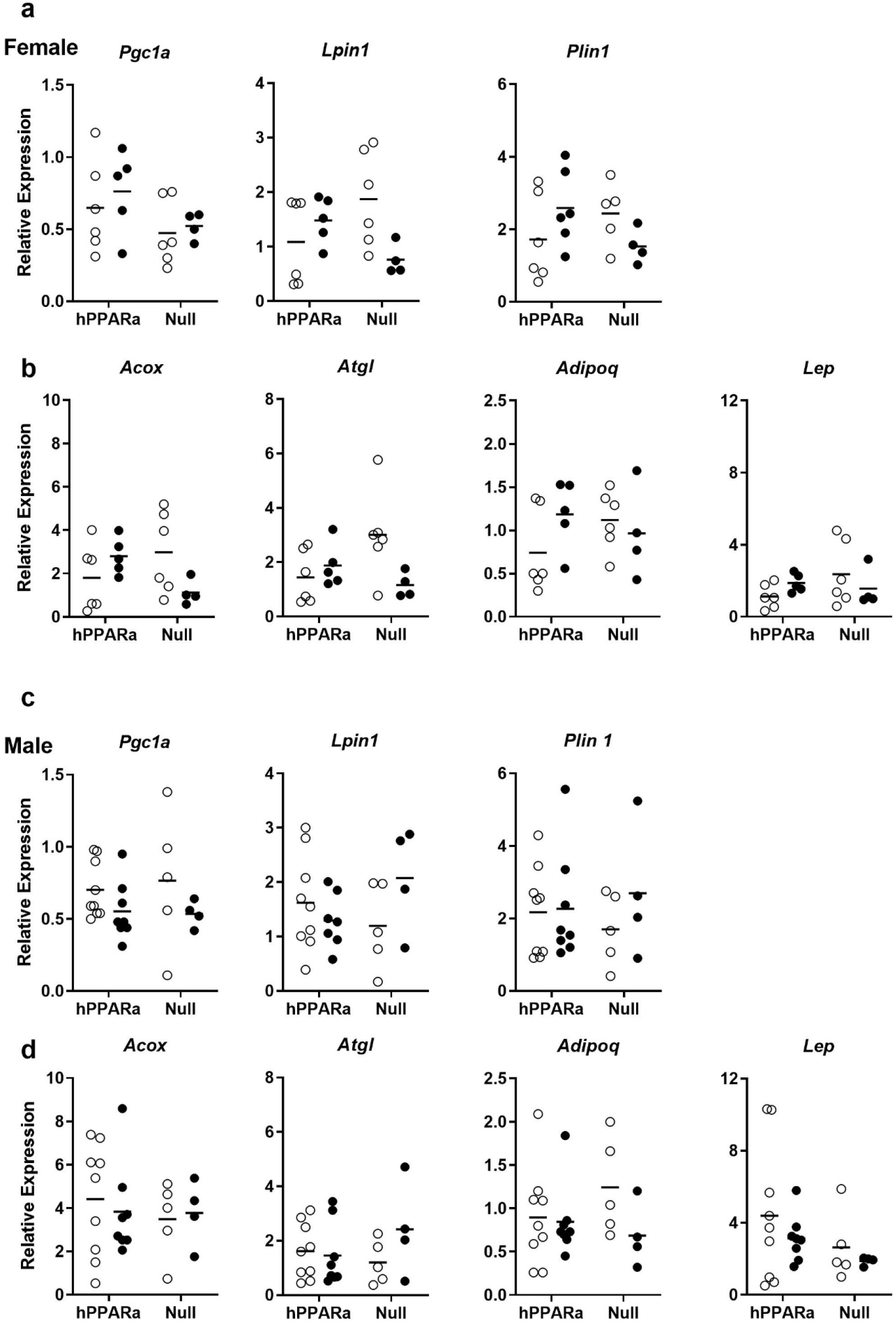
PFOA does not change gene expression related to regulation of fatty acid and triacylglyceride homeostasis in adipose. hPPARα and PPARα null mice were exposed to Vh or PFOA in their drinking water for 6 weeks, as described in Fig. 1. Following isolation of RNA from liver, gene expression was determined by RT-qPCR. **a, c** Transcriptional regulators. **b, d** Functional gene expression. Data are from individual mice, with the mean indicated by a line. N = 5-9.

## Discussion

Exposure to some PFAS has been associated with a number of changes in lipid metabolism, with positive associations between PFAS and lipids reported both in human and animal studies. Most studies have focused on the clinical lipids, and positive associations have been reported particularly with PFOA and PFOS levels and total cholesterol and LDL-C (reviewed in (ATSDR, 2018; EFSA, 2018) and recent studies (Dong et al., 2019; Graber et al., 2019; He et al., 2018; Jain and Ducatman, 2019; Li et al., 2020; C.-Y. Lin et al., 2020; Liu et al., 2020; Seo et al., 2018)), but also with triacylglycerides (Canova et al., 2020; Gardener et al., 2020; Jain and Ducatman, 2019; C. Y. Lin et al., 2020; Seo et al., 2018; Steenland et al., 2010, 2009; Zeng et al., 2015). Studies on the effect of PFAS on lipid composition are just emerging (Dale et al., 2020; Kim et al., 2020; McGlinchey et al., 2020; Pfohl et al., 2020). We generated a more human relevant model, hPPARα mice fed an American diet, in order to test effects of and mechanisms by which PFAS modulate systemic lipid homeostasis (Schlezinger et al., 2020). Here, we tested the hypothesis that PFOA exposure induces liver and serum dyslipidemia in mice expressing hPPARα, in a PPARα-dependent manner. PFOA modulated multiple TG classes in liver and serum, in a sex- and genotype-dependent manner, largely through modification of gene expression in liver but not adipose. The effect of PFOA on serum TG was strongest in female, hPPARα null mice, with PFOA increasing serum TG concentrations by 1.7-fold. The data suggest that effects of PFOA on TG homeostasis are sex- and genotype-dependent and that MIE(s) beyond PPARα play a critical role in PFAS-induced changes in serum TG. When these data are placed in the context of the literature, as discussed below, it becomes clear that human relevant body burdens studies in mice recapitulate the positive association between PFAS and serum TG documented in epidemiological studies.

We previously reported that both the hPPARα female and male mice exposed to PFOA, fed an American diet developed hepatomegaly, associated with histologically evident increases in hepatocyte lipid accumulation (Schlezinger et al., 2020). As triacylglycerides are key for fatty acid storage and accumulation, here, we show that liver:body weight is highly associated with PFOA-induced TG accumulation in hPPARα mice. This is entirely consistent with other researchers who have shown that PFOA induced hepatic TG accumulation occurs to a greater extent in male hPPARα than mPPARα-expressing mice in a similar 6 week exposure scenario (female mice were not studied)(Nakagawa et al., 2012). Quantification of biomarker genes representing the multiple steps and processes involved in hepatic lipid homeostasis showed that genes involved in fatty acid uptake, activation and esterification were strongly upregulated in an hPPARα-dependent manner in both sexes. Induction of these genes is consistent with an accumulation of TG in the liver. However, PFOA also induced genes involved in genes involved in lypolysis and fatty acid oxidation (both peroxisomal and mitochondrial), in an hPPARα-dependent manner. Induction of lipolytic and oxidative enzyme gene expression would be expected to decrease liver TG. However, gene expression does not necessarily reflect the balance of activities of enzymes in the different processes. In previous work in which PFAS induced hepatic TG accumulation it was argued that the balance of fatty acid/TG synthesis and oxidation is skewed toward synthesis by PFAS (Das et al., 2017).

Lipidome analyses of the liver revealed that PFOA exposure not only increased TG abundance, but also modified the composition of the individual TGs. PFOA significantly increased the abundance of MUFA-TG and SFA-TG in hPPARα female and male mice, and these increases were dependent upon hPPARα. SFA are toxic to hepatocytes (S. Nakamura et al., 2009), and cells defend against the lipotoxic effects of SFA by incorporating them into triacylglycerides (Listenberger et al., 2003). MUFA are more readily incorporated into TG than SFAs, therefore desaturase activity is essential for protection against lipotoxicity (Listenberger et al., 2003). Interestingly, studies in rats showed that while peroxisomal beta-oxidation activity remained strongly elevated through a 26 week exposure to PFOA, stearoyl-CoA desaturase activity was only transiently elevated (Uy-Yu et al., 1990). Additional studies will be needed to assess the free fatty acid composition in liver of PFOA-exposed hPPARα mice, as well as desaturase activity, to determine if PFOA stimulates greater SFA accumulation relative to MUFA accumulation.

In the American diet-fed mice, lack of PPARα expression alone increased hepatic TG accumulation relative to hPPARα mice, including TGs substituted will all types of fatty acids, regardless of sex. Hepatic-specific and full body PPARα knockout reduces fatty acid catabolism (lipolysis and oxidation), resulting in TG accumulation and steatosis in the liver following fasting (Montagner et al., 2016; Selen et al., 2021) or consumption of a high fat diet (Régnier et al., 2020). PFOA did not increase liver TGs in the PPARα null mice. This is in contrast to other studies, which did show an increase in liver TGs stimulated by PFOA in PPARα mice (Das et al., 2017; Nakagawa et al., 2012; T. Nakamura et al., 2009). The most likely explanation for the discrepancy is differences in diet, as the previous studies used standard rodent diets. High fat diet feeding of PPARα null mice substantially increases hepatic triacylglycerides (Régnier et al., 2020), to a level that may be what is physiologically-maximal.

In contrast to the observations in liver, PFOA had little effect on serum TG concentrations in serum of hPPARα mice. This is in line with the observation that genes involved in TG export (i.e., very low-density lipoprotein assembly) were either only modestly induced (i.e., *Mttp* induction was only revealed in the combined data in the MLR) or not affected by PFOA (*Apob*). A single previous study has reported the effects of PFAS on lipid homeostasis in female mice; however, serum TGs were not quantified in that study (Rebholz et al., 2016). In males, contradictory data have been reported on the effect of PFOA on serum triacylglycerides, with many studies reporting PFOA-induced decreases in serum TGs (As reviewed in (EFSA, 2018)). We found two other studies in the literature that conducted PFOA exposures over a similar time period as our study (4-6 weeks) and that reported both serum PFOA and TG concentrations (Pouwer et al., 2019; Yan et al., 2014). Across the three studies, at lower PFOA body burdens, increased serum triacylglycerides are observed (**Fig. 12**). At high PFOA body burdens (above those measured even in fluorochemical workers in the US), decreased serum triacylglycerides are observed (**Fig. 12**). A similar non-monotonic dose response was reported in a PFOA exposure study conducted over a two week time period, as well (Loveless et al., 2006). Interestingly, the PFOA serum concentration in this study (≈ 50 ug/ml) fell near the apparent “null point” of the non-monotonic response at which the effect of PFOA changes from increase serum TG to decreasing serum TG. To induce a non-monotonic dose response, at least two different MIEs are necessary (Conolly and Lutz, 2004), with one acting at low concentrations and a second acting at high concentrations. Potential MIEs contributing to PFOA’s effect on serum TGs are discussed below.

**Figure 12.**
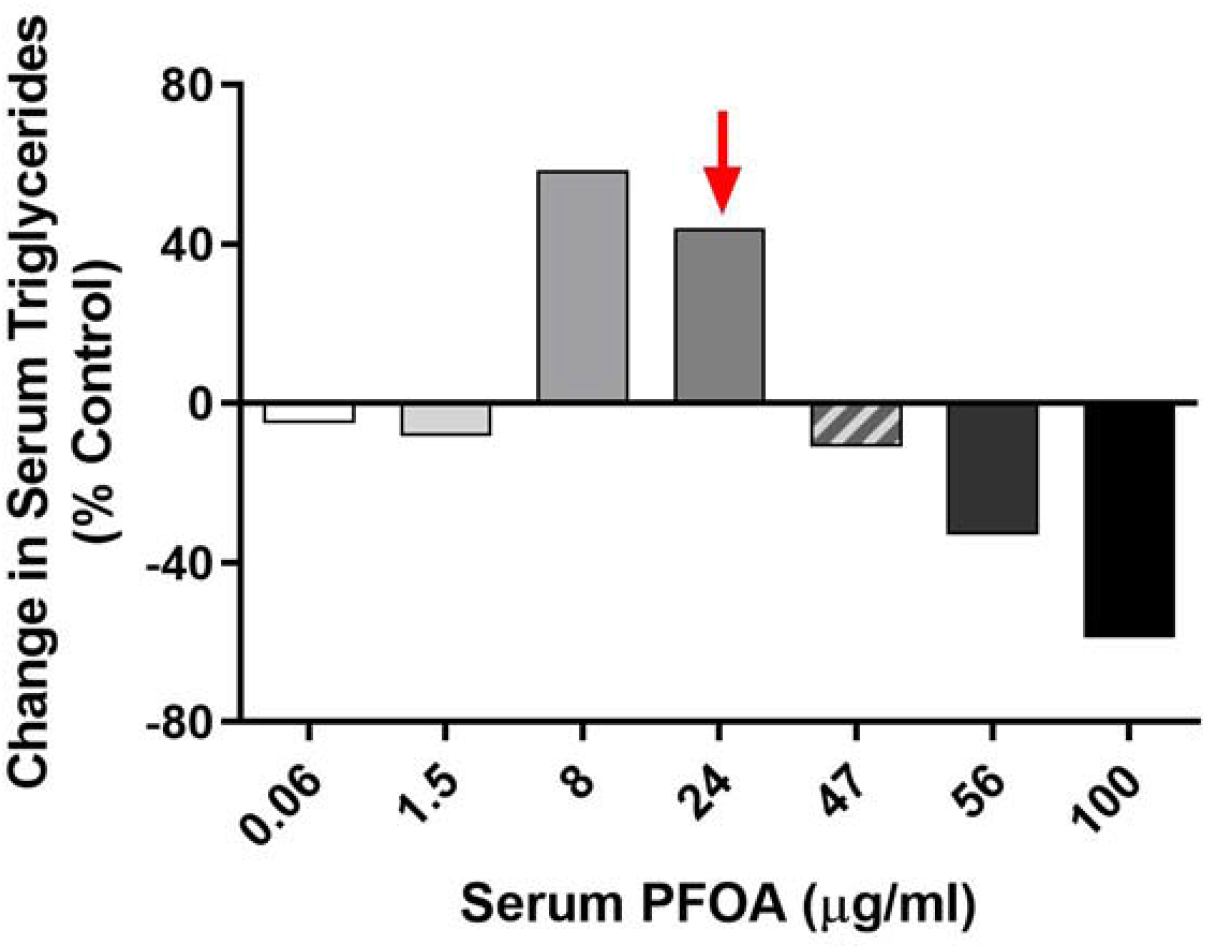
PFOA body burden-dependent changes in serum triacylglycerides. Data were collated across this study (hatched bar), as well as (Yan et al., 2014) and (Pouwer et al., 2019) (solid bars) in which serum TG and PFOA were reported. Data are from males only. Red arrow indicates the PFOA serum level in highly exposed workers.

Intriguingly, PFOA induced a significant increase in serum TGs in PPARα null, females (with a similar trend in males). Previous studies have shown that in PPARα knockout mice there is an increase in vLDL and in serum, with a stronger effect in females than males (Lindén et al., 2001; Stec et al., 2019). TGs associated with vLDL were 2-fold higher in PPARα null than hPPARα females. Total serum TG and vLDL-associated TG were significantly increased by PFOA in PPARα null mice. A similar pattern was seen in males, but the increases were smaller. These data show definitively that PFOA can induce hypertriacylglyceridemia.

The biomarker gene analysis did not reveal a potential mechanism to explain the strong increase in serum TG in female PPARα null mice. However, we do have some clues as to regulatory pathways beyond PPARα and differences between males and females. First, constitutive androstane receptor (CAR) gene targets are strongly upregulated in the livers of these mice (Schlezinger et al., 2020). CAR activation significantly decreases hepatic expression of fatty acid oxidation genes and thus increases serum TG levels (Chen et al., 2019; Maglich et al., 2009; Rezen et al., 2009). Thus, when competition with hPPARα is removed in the PPARα null mice, the positive effect of CAR on serum TG levels predominated. Second, there were significant sex differences in the expression of two transcriptional regulators involved in serum lipid homeostasis, *Pgc1a* and *Lpin1*, which were downregulated by PFOA only in female mice and in an hPPARα-independent manner. PGC1α plays an important role in energy homeostasis (Lin et al., 2004). Lipin 1 interacts with PGC1α to upregulate fatty acid oxidation, mitochondrial oxidative phosphorylation and decrease lipogenesis; loss of lipin 1 increases serum triacylglycerides (Finck et al., 2006). Further research is needed to identify molecular initiating events triggered by PFAS that contribute to regulation of serum lipids. We hypothesize that one will predominate at lower PFAS body burdens and at least one other will predominate at higher PFAS body burdens, generating the non-monotonic dose response we described above.

Atherogenic dyslipidemia (elevated triacylglycerides, decreased HDL-C) and increased blood LDL-C are major contributors to cardiovascular disease (Yusuf et al., 2004), the leading cause of mortality in the US (GBD2015, 2016). Data are accumulating showing a positive association between PFAS and cardiovascular disease risk (De Toni et al., 2020; Huang et al., 2018; Lin et al., 2013; Shankar et al., 2012). Previously we showed that PFOA modulates expression of genes involved in regulating cholesterol homeostasis in liver and increases serum LDL-C (Schlezinger et al., 2020). Here, we show that PFOA modifies both the liver and serum TG in hPPARα-dependent and -independent and in sex-dependent manners. Integration of these data with those in the literature show that lower PFOA body burdens may be more detrimental to lipid homeostasis than high, non-human relevant body burdens. Thus, further research is needed to examine dose response relationships in a human relevant model, as well as to dissect the mechanisms of action leading to disruption of lipid homeostasis by PFAS in females and males.

## Acknowledgements

The authors thank Mr. Nathan Burritt and Daniel Duberg for their excellent technical assistance.

## References

Alves-Bezerra, M., Cohen, D.E., 2017. Triglyceride Metabolism in the Liver. Compr. Physiol. 8, 1–8. https://doi.org/10.1002/cphy.c170012

Appleman, T.D., Higgins, C.P., Quinones, O., Vanderford, B.J., Kolstad, C., Zeigler-Holady, J.C., Dickenson, E.R., 2014. Treatment of poly- and perfluoroalkyl substances in U.S. full-scale water treatment systems. Water Res 51, 246–255. https://doi.org/10.1016/j.watres.2013.10.067

ATSDR, 2018. Toxicological Profile of Perfluoroalkyls [WWW Document]. URL https://www.atsdr.cdc.gov/toxprofiles/tp.asp?id=1117&tid=237 (accessed 2.6.20).

Bjork, J.A., Butenhoff, J.L., Wallace, K.B., 2011. Multiplicity of nuclear receptor activation by PFOA and PFOS in primary human and rodent hepatocytes. Toxicology 288, 8–17. https://doi.org/10.1016/j.tox.2011.06.012

Buhrke, T., Kibellus, A., Lampen, A., 2013. In vitro toxicological characterization of perfluorinated carboxylic acids with different carbon chain lengths. Toxicol. Lett. 218, 97– 104. https://doi.org/10.1016/j.toxlet.2013.01.025

Buhrke, T., Kruger, E., Pevny, S., Rossler, M., Bitter, K., Lampen, A., 2015. Perfluorooctanoic acid (PFOA) affects distinct molecular signalling pathways in human primary hepatocytes. Toxicology 333, 53–62. https://doi.org/10.1016/j.tox.2015.04.004

Calderon-Dominguez, M., Mir, J.F., Fucho, R., Weber, M., Serra, D., Herrero, L., 2016. Fatty acid metabolism and the basis of brown adipose tissue function. Adipocyte 5, 98–118. https://doi.org/10.1080/21623945.2015.1122857

Canova, C., Barbieri, G., Zare Jeddi, M., Gion, M., Fabricio, A., Daprà, F., Russo, F., Fletcher, T., Pitter, G., 2020. Associations between perfluoroalkyl substances and lipid profile in a highly exposed young adult population in the Veneto Region.Environ. Int. 145, 106117. https://doi.org/10.1016/j.envint.2020.106117

CDC, 2019. Fourth National Report on Human Exposure to Environmental Chemicals. Updated Tables, January 2019.

Chen, F., Coslo, D.M., Chen, T., Zhang, L., Tian, Y., Smith, P.B., Patterson, A.D., Omiecinski, C.J., 2019. Metabolomic Approaches Reveal the Role of CAR in Energy Metabolism. J. Proteome Res. 18, 239–251. https://doi.org/10.1021/acs.jproteome.8b00566

Chen, Z., Gropler, M.C., Norris, J., Lawrence, J.C.J., Harris, T.E., Finck, B.N., 2008. Alterations in hepatic metabolism in fld mice reveal a role for lipin 1 in regulating VLDL-triacylglyceride secretion. Arterioscler. Thromb. Vasc. Biol. 28, 1738–1744. https://doi.org/10.1161/ATVBAHA.108.171538

Conolly, R.B., Lutz, W.K., 2004. Nonmonotonic dose-response relationships: mechanistic basis, kinetic modeling, and implications for risk assessment. Toxicol. Sci. 77, 151–157. https://doi.org/10.1093/toxsci/kfh007

Cousins, I.T., Vestergren, R., Wang, Z., Scheringer, M., McLachlan, M.S., 2016. The precautionary principle and chemicals management: The example of perfluoroalkyl acids in groundwater. Environ. Int. 94, 331–340. https://doi.org/10.1016/j.envint.2016.04.044

Dale, K., Yadetie, F., Müller, M.B., Pampanin, D.M., Gilabert, A., Zhang, X., Tairova, Z., Haarr, A., Lille-Langøy, R., Lyche, J.L., Porte, C., Karlsen, O.A., Goksøyr, A., 2020. Proteomics and lipidomics analyses reveal modulation of lipid metabolism by perfluoroalkyl substances in liver of Atlantic cod (Gadus morhua). Aquat. Toxicol. 227, 105590. https://doi.org/10.1016/j.aquatox.2020.105590

Das, K.P., Wood, C.R., Lin, M.J., Starkov, A.A., Lau, C., Wallace, K.B., Corton, J.C., Abbott, B.D., 2017. Perfluoroalkyl acids-induced liver steatosis: Effects on genes controlling lipid homeostasis. Toxicology 378, 37–52. https://doi.org/10.1016/j.tox.2016.12.007

De Toni, L., Radu, C.M., Sabovic, I., Di Nisio, A., Dall’Acqua, S., Guidolin, D., Spampinato, S., Campello, E., Simioni, P., Foresta, C., 2020. Increased Cardiovascular Risk Associated with Chemical Sensitivity to Perfluoro-Octanoic Acid: Role of Impaired Platelet Aggregation. Int. J. Mol. Sci. 21. https://doi.org/10.3390/ijms21020399

Dong, Z., Wang, H., Yu, Y.Y., Li, Y.B., Naidu, R., Liu, Y., 2019. Using 2003-2014 U.S. NHANES data to determine the associations between per-and polyfluoroalkyl substances and cholesterol: Trend and implications. Ecotoxicol. Environ. Saf. 173, 461–468. https://doi.org/10.1016/j.ecoenv.2019.02.061

EFSA, 2018. Risk to human health related to the presence of perfluorooctane sulfonic acid and perfluorooctanoic acid in food. EFSA J 16, 5194.

Finck, B.N., Gropler, M.C., Chen, Z., Leone, T.C., Croce, M.A., Harris, T.E., Lawrence, J.C.J., Kelly, D.P., 2006. Lipin 1 is an inducible amplifier of the hepatic PGC-1alpha/PPARalpha regulatory pathway. Cell Metab. 4, 199–210. https://doi.org/10.1016/j.cmet.2006.08.005

Fruchart, J.-C., Duriez, P., 2006. Mode of action of fibrates in the regulation of triglyceride and HDL-cholesterol metabolism. Drugs Today (Barc). 42, 39–64. https://doi.org/10.1358/dot.2006.42.1.963528

Frühbeck, G., Catalán, V., Rodríguez, A., Gómez-Ambrosi, J., 2018. Adiponectin-leptin ratio: A promising index to estimate adipose tissue dysfunction. Relation with obesity-associated cardiometabolic risk. Adipocyte 7, 57–62. https://doi.org/10.1080/21623945.2017.1402151

Galarraga, M., Campion, J., Munoz-Barrutia, A., Boque, N., Moreno, H., Martinez, J.A., Milagro, F., Ortiz-de-Solorzano, C., 2012. Adiposoft: automated software for the analysis of white adipose tissue cellularity in histological sections. J Lipid Res 53, 2791–2796. https://doi.org/10.1194/jlr.D023788

Gardener, H., Sun, Q., Grandjean, P., 2020. PFAS concentration during pregnancy in relation to cardiometabolic health and birth outcomes. Environ. Res. 192, 110287. https://doi.org/10.1016/j.envres.2020.110287

GBD2015, 2016. Global, regional, and national life expectancy, all-cause mortality, and cause-specific mortality for 249 causes of death, 1980-2015: a systematic analysis for the Global Burden of Disease Study 2015. Lancet 388, 1459–1544. https://doi.org/10.1016/S0140-6736(16)31012-1

Graber, J.M., Alexander, C., Laumbach, R.J., Black, K., Strickland, P.O., Georgopoulos, P.G., Marshall, E.G., Shendell, D.G., Alderson, D., Mi, Z., Mascari, M., Weisel, C.P., 2019. Per and polyfluoroalkyl substances (PFAS) blood levels after contamination of a community water supply and comparison with 2013-2014 NHANES. J. Expo. Sci. Environ. Epidemiol. 29, 172–182. https://doi.org/10.1038/s41370-018-0096-z

Guelfo, J.L., Marlow, T., Klein, D.M., Savitz, D.A., Frickel, S., Crimi, M., Suuberg, E.M., 2018. Evaluation and Management Strategies for Per- and Polyfluoroalkyl Substances (PFASs) in Drinking Water Aquifers: Perspectives from Impacted U.S. Northeast Communities. Environ. Health Perspect. 126, 65001. https://doi.org/10.1289/EHP2727

He, X., Liu, Y., Xu, B., Gu, L., Tang, W., 2018. PFOA is associated with diabetes and metabolic alteration in US men: National Health and Nutrition Examination Survey 2003-2012. Sci. Total Environ. 625, 566–574. https://doi.org/10.1016/j.scitotenv.2017.12.186

Hu, X.C., Andrews, D.Q., Lindstrom, A.B., Bruton, T.A., Schaider, L.A., Grandjean, P., Lohmann, R., Carignan, C.C., Blum, A., Balan, S.A., Higgins, C.P., Sunderland, E.M., 2016. Detection of Poly- and Perfluoroalkyl Substances (PFASs) in U.S. Drinking Water Linked to Industrial Sites, Military Fire Training Areas, and Wastewater Treatment Plants. Environ. Sci. Technol. Lett. 3, 344–350. https://doi.org/10.1021/acs.estlett.6b00260

Huang, M., Jiao, J., Zhuang, P., Chen, X., Wang, J., Zhang, Y., 2018. Serum polyfluoroalkyl chemicals are associated with risk of cardiovascular diseases in national US population. Environ. Int. 119, 37–46. https://doi.org/10.1016/j.envint.2018.05.051

Jain, R.B., Ducatman, A., 2019. Roles of gender and obesity in defining correlations between perfluoroalkyl substances and lipid/lipoproteins. Sci. Total Environ. 653, 74–81. https://doi.org/10.1016/j.scitotenv.2018.10.362

Keller, H., Devchand, P.R., Perroud, M., Wahli, W., 1997. PPAR alpha structure-function relationships derived from species-specific differences in responsiveness to hypolipidemic agents. Biol Chem 378, 651–655. https://doi.org/10.1515/bchm.1997.378.7.651

Kersten, S., Stienstra, R., 2017. The role and regulation of the peroxisome proliferator activated receptor alpha in human liver. Biochimie 136, 75–84. https://doi.org/10.1016/j.biochi.2016.12.019

Kim, D.-H., Lee, J.-H., Oh, J.-E., 2019. Assessment of individual-based perfluoroalkly substances exposure by multiple human exposure sources. J. Hazard. Mater. 365, 26–33. https://doi.org/10.1016/j.jhazmat.2018.10.066

Kim, H.M., Long, N.P., Yoon, S.J., Anh, N.H., Kim, S.J., Park, J.H., Kwon, S.W., 2020. Omics approach reveals perturbation of metabolism and phenotype in Caenorhabditis elegans triggered by perfluorinated compounds. Sci. Total Environ. 703, 135500. https://doi.org/10.1016/j.scitotenv.2019.135500

Li, Y., Barregard, L., Xu, Y., Scott, K., Pineda, D., Lindh, C.H., Jakobsson, K., Fletcher, T., 2020. Associations between perfluoroalkyl substances and serum lipids in a Swedish adult population with contaminated drinking water. Environ. Health 19, 33. https://doi.org/10.1186/s12940-020-00588-9

Lin, C.-Y., Lee, H.-L., Hwang, Y.-T., Su, T.-C., 2020. The association between total serum isomers of per- and polyfluoroalkyl substances, lipid profiles, and the DNA oxidative/nitrative stress biomarkers in middle-aged Taiwanese adults. Environ. Res. 182, 109064. https://doi.org/10.1016/j.envres.2019.109064

Lin, C.-Y., Lin, L.-Y., Wen, T.-W., Lien, G.-W., Chien, K.-L., Hsu, S.H.J., Liao, C.-C., Sung, F.-C., Chen, P.-C., Su, T.-C., 2013. Association between levels of serum perfluorooctane sulfate and carotid artery intima-media thickness in adolescents and young adults. Int. J. Cardiol. 168, 3309–3316. https://doi.org/10.1016/j.ijcard.2013.04.042

Lin, C.Y., Lee, H.L., Hwang, Y.T., Su, T.C., 2020. The association between total serum isomers of per- and polyfluoroalkyl substances, lipid profiles, and the DNA oxidative/nitrative stress biomarkers in middle-aged Taiwanese adults. Environ. Res. 182, 109064. https://doi.org/10.1016/j.envres.2019.109064

Lin, J., Wu, P.-H., Tarr, P.T., Lindenberg, K.S., St-Pierre, J., Zhang, C.-Y., Mootha, V.K., Jäger, S., Vianna, C.R., Reznick, R.M., Cui, L., Manieri, M., Donovan, M.X., Wu, Z., Cooper, M.P., Fan, M.C., Rohas, L.M., Zavacki, A.M., Cinti, S., Shulman, G.I., Lowell, B.B., Krainc, D., Spiegelman, B.M., 2004. Defects in adaptive energy metabolism with CNS-linked hyperactivity in PGC-1alpha null mice. Cell 119, 121–135. https://doi.org/10.1016/j.cell.2004.09.013

Lin, T.-W., Chen, M.-K., Lin, C.-C., Chen, M.-H., Tsai, M.-S., Chan, D.-C., Hung, K.-Y., Chen, P.-C., 2020. Association between exposure to perfluoroalkyl substances and metabolic syndrome and related outcomes among older residents living near a Science Park in Taiwan. Int. J. Hyg. Environ. Health 230, 113607. https://doi.org/10.1016/j.ijheh.2020.113607

Lindén, D., Alsterholm, M., Wennbo, H., Oscarsson, J., 2001. PPARalpha deficiency increases secretion and serum levels of apolipoprotein B-containing lipoproteins. J. Lipid Res. 42, 1831–1840.

Listenberger, L.L., Han, X., Lewis, S.E., Cases, S., Farese, R.V.J., Ory, D.S., Schaffer, J.E., 2003. Triglyceride accumulation protects against fatty acid-induced lipotoxicity. Proc. Natl. Acad. Sci. U. S. A. 100, 3077–3082. https://doi.org/10.1073/pnas.0630588100

Liu, G., Zhang, B., Hu, Y., Rood, J., Liang, L., Qi, L., Bray, G.A., DeJonge, L., Coull, B., Grandjean, P., Furtado, J.D., Sun, Q., 2020. Associations of Perfluoroalkyl substances with blood lipids and Apolipoproteins in lipoprotein subspecies: the POUNDS-lost study. Environ. Health 19, 5. https://doi.org/10.1186/s12940-020-0561-8

Loveless, S.E., Finlay, C., Everds, N.E., Frame, S.R., Gillies, P.J., O’Connor, J.C., Powley, C.R., Kennedy, G.L., 2006. Comparative responses of rats and mice exposed to linear/branched, linear, or branched ammonium perfluorooctanoate (APFO). Toxicology 220, 203–217. https://doi.org/10.1016/j.tox.2006.01.003

Lutkewitte, A.J., McCommis, K.S., Schweitzer, G.G., Chambers, K.T., Graham, M.J., Wang, L., Patti, G.J., Hall, A.M., Finck, B.N., 2019. Hepatic monoacylglycerol acyltransferase 1 is induced by prolonged food deprivation to modulate the hepatic fasting response. J. Lipid Res. 60, 528–538. https://doi.org/10.1194/jlr.M089722

Maglich, J.M., Lobe, D.C., Moore, J.T., 2009. The nuclear receptor CAR (NR1I3) regulates serum triglyceride levels under conditions of metabolic stress. J. Lipid Res. 50, 439–445. https://doi.org/10.1194/jlr.M800226-JLR200

Makey, C.M., Webster, T.F., Martin, J.W., Shoeib, M., Harner, T., Dix-Cooper, L., Webster, G.M., 2017. Airborne Precursors Predict Maternal Serum Perfluoroalkyl Acid Concentrations. Environ. Sci. Technol. 51, 7667–7675. https://doi.org/10.1021/acs.est.7b00615

Maloney, E. k., Waxman, D.J., 1999. trans-Activation of PPARa and PPARg by structurally diverse environmental chemicals. Toxicol. Appl. Pharmacol. 161, 209–218.

Matilla-Santander, N., Valvi, D., Lopez-Espinosa, M.J., Manzano-Salgado, C.B., Ballester, F., Ibarluzea, J., Santa-Marina, L., Schettgen, T., Guxens, M., Sunyer, J., Vrijheid, M., 2017. Exposure to Perfluoroalkyl Substances and Metabolic Outcomes in Pregnant Women: Evidence from the Spanish INMA Birth Cohorts. Env. Heal. Perspect 125, 117004. https://doi.org/10.1289/EHP1062

McGlinchey, A., Sinioja, T., Lamichhane, S., Sen, P., Bodin, J., Siljander, H., Dickens, A.M., Geng, D., Carlsson, C., Duberg, D., Ilonen, J., Virtanen, S.M., Dirven, H., Berntsen, H.F., Zimmer, K., Nygaard, U.C., Orešič, M., Knip, M., Hyötyläinen, T., 2020. Prenatal exposure to perfluoroalkyl substances modulates neonatal serum phospholipids, increasing risk of type 1 diabetes. Environ. Int. 143, 105935. https://doi.org/10.1016/j.envint.2020.105935

Minata, M., Harada, K.H., Karrman, A., Hitomi, T., Hirosawa, M., Murata, M., Gonzalez, F.J., Koizumi, A., 2010. Role of peroxisome proliferator-activated receptor-alpha in hepatobiliary injury induced by ammonium perfluorooctanoate in mouse liver. Ind Heal. 48, 96–107. https://doi.org/10.2486/indhealth.48.96

Montagner, A., Polizzi, A., Fouché, E., Ducheix, S., Lippi, Y., Lasserre, F., Barquissau, V., Régnier, M., Lukowicz, C., Benhamed, F., Iroz, A., Bertrand-Michel, J., Al Saati, T., Cano, P., Mselli-Lakhal, L., Mithieux, G., Rajas, F., Lagarrigue, S., Pineau, T., Loiseau, N., Postic, C., Langin, D., Wahli, W., Guillou, H., 2016. Liver PPARα is crucial for whole-body fatty acid homeostasis and is protective against NAFLD. Gut 65, 1202–1214. https://doi.org/10.1136/gutjnl-2015-310798

Morimura, K., Cheung, C., Ward, J.M., Reddy, J.K., Gonzalez, F.J., 2006. Differential susceptibility of mice humanized for peroxisome proliferator-activated receptor alpha to Wy-14,643-induced liver tumorigenesis. Carcinogenesis 27, 1074–1080. https://doi.org/10.1093/carcin/bgi329

Nadra, K., Médard, J.-J., Mul, J.D., Han, G.-S., Grès, S., Pende, M., Metzger, D., Chambon, P., Cuppen, E., Saulnier-Blache, J.-S., Carman, G.M., Desvergne, B., Chrast, R., 2012. Cell autonomous lipin 1 function is essential for development and maintenance of white and brown adipose tissue. Mol. Cell. Biol. 32, 4794–4810. https://doi.org/10.1128/MCB.00512-12

Nakagawa, T., Ramdhan, D.H., Tanaka, N., Naito, H., Tamada, H., Ito, Y., Li, Y., Hayashi, Y., Yamagishi, N., Yanagiba, Y., Aoyama, T., Gonzalez, F.J., Nakajima, T., 2012. Modulation of ammonium perfluorooctanoate-induced hepatic damage by genetically different PPARα in mice. Arch. Toxicol. 86, 63–74. https://doi.org/10.1007/s00204-011-0704-3

Nakamura, S., Takamura, T., Matsuzawa-Nagata, N., Takayama, H., Misu, H., Noda, H., Nabemoto, S., Kurita, S., Ota, T., Ando, H., Miyamoto, K.-I., Kaneko, S., 2009. Palmitate induces insulin resistance in H4IIEC3 hepatocytes through reactive oxygen species produced by mitochondria. J. Biol. Chem. 284, 14809–14818. https://doi.org/10.1074/jbc.M901488200

Nakamura, T., Ito, Y., Yanagiba, Y., Ramdhan, D.H., Kono, Y., Naito, H., Hayashi, Y., Li, Y., Aoyama, T., Gonzalez, F.J., Nakajima, T., 2009. Microgram-order ammonium perfluorooctanoate may activate mouse peroxisome proliferator-activated receptor alpha, but not human PPARalpha. Toxicology 265, 27–33. https://doi.org/10.1016/j.tox.2009.09.004

Norris, A.W., Chen, L., Fisher, S.J., Szanto, I., Ristow, M., Jozsi, A.C., Hirshman, M.F., Rosen, E.D., Goodyear, L.J., Gonzalez, F.J., Spiegelman, B.M., Kahn, C.R., 2003. Muscle-specific PPARgamma-deficient mice develop increased adiposity and insulin resistance but respond to thiazolidinediones. J Clin Invest 112, 608–618. https://doi.org/10.1172/JCI17305

Oswal, D.P., Alter, G.M., Rider Jr., S.D., Hostetler, H.A., 2014. A single amino acid change humanizes long-chain fatty acid binding and activation of mouse peroxisome proliferator-activated receptor alpha. J Mol Graph Model 51, 27–36. https://doi.org/10.1016/j.jmgm.2014.04.006

Pang, Z., Chong, J., Li, S., Xia, J., 2020. MetaboAnalystR 3.0: Toward an Optimized Workflow for Global Metabolomics. Metabolites 10. https://doi.org/10.3390/metabo10050186

Peng, S., Yan, L., Zhang, J., Wang, Z., Tian, M., Shen, H., 2013. An integrated metabonomics and transcriptomics approach to understanding metabolic pathway disturbance induced by perfluorooctanoic acid. J Pharm Biomed Anal 86, 56–64. https://doi.org/10.1016/j.jpba.2013.07.014

Pfaffl, M.W., 2001. A new mathematical model for relative quantification in real-time RT-PCR. Nucleic Acids Res 29, e45.

Pfohl, M., Ingram, L., Marques, E., Auclair, A., Barlock, B., Jamwal, R., Anderson, D., Cummings, B.S., Slitt, A.L., 2020. Perfluorooctanesulfonic Acid and Perfluorohexanesulfonic Acid Alter the Blood Lipidome and the Hepatic Proteome in a Murine Model of Diet-Induced Obesity. Toxicol. Sci. 178, 311–324. https://doi.org/10.1093/toxsci/kfaa148

Pouwer, M.G., Pieterman, E.J., Chang, S.C., Olsen, G.W., Caspers, M.P.M., Verschuren, L., Jukema, J.W., Princen, H.M.G., 2019. Dose Effects of Ammonium Perfluorooctanoate on Lipoprotein Metabolism in APOE*3-Leiden.CETP Mice. Toxicol Sci 168, 519–534. https://doi.org/10.1093/toxsci/kfz015

Puigserver, P., Wu, Z., Park, C.W., Graves, R., Wright, M., Spiegelman, B.M., 1998. A cold-inducible coactivator of nuclear receptors linked to adaptive thermogenesis. Cell 92, 829– 839.

Rakhshandehroo, M., Hooiveld, G., Muller, M., Kersten, S., 2009. Comparative analysis of gene regulation by the transcription factor PPARalpha between mouse and human. PLoS One 4, e6796. https://doi.org/10.1371/journal.pone.0006796

Rebholz, S.L., Jones, T., Herrick, R.L., Xie, C., Calafat, A.M., Pinney, S.M., Woollett, L.A., 2016. Hypercholesterolemia with consumption of PFOA-laced Western diets is dependent on strain and sex of mice. Toxicol Rep 3, 46–54. https://doi.org/10.1016/j.toxrep.2015.11.004

Régnier, M., Polizzi, A., Smati, S., Lukowicz, C., Fougerat, A., Lippi, Y., Fouché, E., Lasserre, F., Naylies, C., Bétoulières, C., Barquissau, V., Mouisel, E., Bertrand-Michel, J., Batut, A., Saati, T.Al, Canlet, C., Tremblay-Franco, M., Ellero-Simatos, S., Langin, D., Postic, C., Wahli, W., Loiseau, N., Guillou, H., Montagner, A., 2020. Hepatocyte-specific deletion of Pparα promotes NAFLD in the context of obesity. Sci. Rep. 10, 6489. https://doi.org/10.1038/s41598-020-63579-3

Rezen, T., Tamasi, V., Lövgren-Sandblom, A., Björkhem, I., Meyer, U.A., Rozman, D., 2009. Effect of CAR activation on selected metabolic pathways in normal and hyperlipidemic mouse livers. BMC Genomics 10, 384. https://doi.org/10.1186/1471-2164-10-384

Rosen, M.B., Das, K.P., Rooney, J., Abbott, B., Lau, C., Corton, J.C., 2017. PPARalpha-independent transcriptional targets of perfluoroalkyl acids revealed by transcript profiling. Toxicology 387, 95–107. https://doi.org/10.1016/j.tox.2017.05.013

Rosenmai, A.K., Ahrens, L., le Godec, T., Lundqvist, J., Oskarsson, A., 2018. Relationship between peroxisome proliferator-activated receptor alpha activity and cellular concentration of 14 perfluoroalkyl substances in HepG2 cells. J Appl Toxicol 38, 219–226. https://doi.org/10.1002/jat.3515

Rosenwald, M., Efthymiou, V., Opitz, L., Wolfrum, C., 2017. SRF and MKL1 Independently Inhibit Brown Adipogenesis. PLoS One 12, e0170643. https://doi.org/10.1371/journal.pone.0170643

Scheringer, M., Trier, X., Cousins, I.T., de Voogt, P., Fletcher, T., Wang, Z., Webster, T.F., 2014. Helsingør statement on poly- and perfluorinated alkyl substances (PFASs). Chemosphere 114, 337–339. https://doi.org/10.1016/j.chemosphere.2014.05.044

Schlezinger, J., Puckett, H., Nielsen, G., Oliver, J., Heiger-Bernays, W., Webster, T., 2020. Perfluorooctanoic acid activates multiple nuclear receptor pathways and skews expression of genes regulating cholesterol homeostasis in liver of humanized PPARα mice fed an American diet. Toxicol Appl Pharmacol 405, 115204.

Selen, E.S., Choi, J., Wolfgang, M.J., 2021. Discordant hepatic fatty acid oxidation and triglyceride hydrolysis leads to liver disease. JCI insight 6. https://doi.org/10.1172/jci.insight.135626

Seo, S.-H., Son, M.-H., Choi, S.-D., Lee, D.-H., Chang, Y.-S., 2018. Influence of exposure to perfluoroalkyl substances (PFASs) on the Korean general population: 10-year trend and health effects. Environ. Int. 113, 149–161. https://doi.org/10.1016/j.envint.2018.01.025

Shankar, A., Xiao, J., Ducatman, A., 2012. Perfluorooctanoic acid and cardiovascular disease in US adults. Arch. Intern. Med. 172, 1397–1403. https://doi.org/10.1001/archinternmed.2012.3393

Staels, B., Dallongeville, J., Auwerx, J., Schoonjans, K., Leitersdorf, E., Fruchart, J.C., 1998. Mechanism of action of fibrates on lipid and lipoprotein metabolism. Circulation 98, 2088– 2093.

Stec, D.E., Gordon, D.M., Hipp, J.A., Hong, S., Mitchell, Z.L., Franco, N.R., Robison, J.W., Anderson, C.D., Stec, D.F., Hinds, T.D.J., 2019. Loss of hepatic PPARα promotes inflammation and serum hyperlipidemia in diet-induced obesity. Am. J. Physiol. Regul. Integr. Comp. Physiol. 317, R733–R745. https://doi.org/10.1152/ajpregu.00153.2019

Steenland, K., Fletcher, T., Savitz, D.A., 2010. Epidemiologic evidence on the health effects of perfluorooctanoic acid (PFOA). Environ. Health Perspect. 118, 1100–1108. https://doi.org/10.1289/ehp.0901827

Steenland, K., Tinker, S., Frisbee, S., Ducatman, A., Vaccarino, V., 2009. Association of perfluorooctanoic acid and perfluorooctane sulfonate with serum lipids among adults living near a chemical plant. Am. J. Epidemiol. 170, 1268–1278. https://doi.org/10.1093/aje/kwp279

Sundborn, G., Thornley, S., Merriman, T.R., Lang, B., King, C., Lanaspa, M.A., Johnson, R.J., 2019. Are Liquid Sugars Different from Solid Sugar in Their Ability to Cause Metabolic Syndrome? Obesity (Silver Spring). 27, 879–887. https://doi.org/10.1002/oby.22472

Takacs, M.L., Abbott, B.D., 2007. Activation of mouse and human peroxisome proliferator-activated receptors (alpha, beta/delta, gamma) by perfluorooctanoic acid and perfluorooctane sulfonate. Toxicol Sci 95, 108–117. https://doi.org/10.1093/toxsci/kfl135

Tan, X., Xie, G., Sun, X., Li, Q., Zhong, W., Qiao, P., Sun, X., Jia, W., Zhou, Z., 2013. High fat diet feeding exaggerates perfluorooctanoic acid-induced liver injury in mice via modulating multiple metabolic pathways. PLoS One 8, e61409. https://doi.org/10.1371/journal.pone.0061409

USDA, 2018. What we eat in America [WWW Document]. URL https://www.ars.usda.gov/northeast-area/beltsville-md-bhnrc/beltsville-human-nutrition-research-center/food-surveys-research-group/docs/wweianhanes-overview/

Uy-Yu, N., Kawashima, Y., Horii, S., Kozuka, H., 1990. Effects of chronic administration of perfluorooctanoic acid on fatty acid metabolism in rat liver: relationship among stearoyl-coenzyme A desaturase, 1-acylglycerophosphocholine acyltransferase, and acyl composition of microsomal phosphatidylcholine. J. Pharmacobiodyn. 13, 581–590. https://doi.org/10.1248/bpb1978.13.581

Vanden Heuvel, J.P., Thompson, J.T., Frame, S.R., Gillies, P.J., 2006. Differential activation of nuclear receptors by perfluorinated fatty acid analogs and natural fatty acids: a comparison of human, mouse, and rat peroxisome proliferator-activated receptor-alpha, -beta, and - gamma, liver X receptor-beta, and retinoid X rec. Toxicol Sci 92, 476–489. https://doi.org/10.1093/toxsci/kfl014

Vega, R.B., Huss, J.M., Kelly, D.P., 2000. The coactivator PGC-1 cooperates with peroxisome proliferator-activated receptor alpha in transcriptional control of nuclear genes encoding mitochondrial fatty acid oxidation enzymes. Mol. Cell. Biol. 20, 1868–1876. https://doi.org/10.1128/mcb.20.5.1868-1876.2000

Wolf, C.J., Schmid, J.E., Lau, C., Abbott, B.D., 2012. Activation of mouse and human peroxisome proliferator-activated receptor-alpha (PPARalpha) by perfluoroalkyl acids (PFAAs): further investigation of C4-C12 compounds. Reprod Toxicol 33, 546–551. https://doi.org/10.1016/j.reprotox.2011.09.009

Wolf, C.J., Takacs, M.L., Schmid, J.E., Lau, C., Abbott, B.D., 2008. Activation of mouse and human peroxisome proliferator-activated receptor alpha by perfluoroalkyl acids of different functional groups and chain lengths. Toxicol Sci 106, 162–171. https://doi.org/10.1093/toxsci/kfn166

Yan, S., Wang, J., Zhang, W., Dai, J., 2014. Circulating microRNA profiles altered in mice after 28 d exposure to perfluorooctanoic acid. Toxicol Lett 224, 24–31.

Yang, Q., Nagano, T., Shah, Y., Cheung, C., Ito, S., Gonzalez, F.J., 2008. The PPAR alpha-humanized mouse: a model to investigate species differences in liver toxicity mediated by PPAR alpha. Toxicol. Sci. 101, 132–139. https://doi.org/10.1093/toxsci/kfm206

Yusuf, S., Hawken, S., Ounpuu, S., Dans, T., Avezum, A., Lanas, F., McQueen, M., Budaj, A., Pais, P., Varigos, J., Lisheng, L., Investigators, I.S., 2004. Effect of potentially modifiable risk factors associated with myocardial infarction in 52 countries (the INTERHEART study): case-control study. Lancet 364, 937–952. https://doi.org/10.1016/S0140-6736(04)17018-9

Zeng, X.W., Qian, Z., Emo, B., Vaughn, M., Bao, J., Qin, X.D., Zhu, Y., Li, J., Lee, Y.L., Dong, G.H., 2015. Association of polyfluoroalkyl chemical exposure with serum lipids in children. Sci Total Env. 512–513, 364–370. https://doi.org/10.1016/j.scitotenv.2015.01.042

