## Supplemental Material for "Perfluorooctanoic Acid Induces Liver and Serum Dyslipidemia in Humanized PPARα Mice fed an American Diet"

### Table of Contents

| | | Page |
|---|---|---|
| Table S1 | Cohort information. | 2 |
| Table S2 | Components of the "What we eat in America" diet (Research Diets D18110214) | 3 |
| Table S3 | Primer sequences for reverse transcriptase qPCR. | 4 |
| Table S4 | Statistical analyses of fold-change in liver triacylglyceride classes | 5 |
| Table S5 | ANOVA liver triacylglycerides | 5 |
| Table S6 | Statistical analyses of fold-change in serum triacylglyceride classes | 5 |
| Table S7 | ANOVA of serum triacylglycerides | 5 |
| Figure S1 | Spleen/body weight and thymus/body weight ratios in PFOA-treated mice. | 6 |
| Figure S2 | Lipidomic analysis of liver. | 7 |
| Figure S3 | Spearman correlation analysis of liver triacylglycerides | 8 |

**Table S1: Cohort information**

| | | *PPARA* | | | | Knockout | | | |
|---|---|---|---|---|---|---|---|---|---|
| | | Vh | | PFOA | | Vh | | PFOA | |
| **Cohort Number** | **Breeders** | **Male Pups** | **Female Pups** | **Male Pups** | **Female Pups** | **Male Pups** | **Female Pups** | **Male Pups** | **Female Pups** |
| 1 | Male 1/ Female 1 | 1 | 2 | | | | | | |
| 2 | Male 2/ Female 2 | 1 | | 1 | 1 | 1 | 1 | | 1 |
| 3 [a] | Male 1/ Female 1 | 1 | 1 | | | 1 | 1 | | |
| 4 | Male 2/ Female 2 | | | 1 | | | | 1 | 1 |
| 5 | Male 1/ Female 1 | | | 1 | 2 | | | 2 | 2 |
| 6 | Male 2/ Female 2 | 2 | 2 | 1 | | 1 | 3 | | |
| 7 | Male 1/ Female 1 | 3 | | | 1 | 1 | | | |
| 8 [b] | Male 2/ Female 2 | | 1 | 1 | 2 | | | | 1 |
| 9 | Male 2/ Female 1 | | | | | 2 | 1 | 2 | 1 |
| 10 | Male 4/ Female 4 | | 1 | 2 | | | | | |
| 11 [c] | Male 2/ Female 3 | 1 | | 1 | 1 | | | 1 | |
| | **Total** | **9** | **7** | **9** | **7** | **6** | **6** | **5** | **6** |

[a]One female failed to thrive (the final body weight was more than 2 standard deviations lower than the average final body weight) and was excluded from data analysis.
[b]One female had total serum lipids 4 standard deviations above the mean and was excluded from data analysis.
[c]One male had a severe eye injury and was euthanized early.

**Table S2: Components of the "What we eat in America" diet (Research Diets D18110214).**

| | **% Grams** | **% kcal** |
|---|---|---|
| Protein | 16 | 14.7 |
| Carbohydrate | 57 | 51.8 |
| Fat | 16 | 33.5 |
| Fiber | 5.4 | |
| Cholesterol | 0.05 | |
| Kcal/gm | 4.4 | |
| **Ingredient** | **grams** | **kcal** |
| Casein | 146 | 584 |
| L-cystine | 3 | 12 |
| Corn Starch | 176.81 | 707 |
| Maltodextrin 10 | 100 | 400 |
| Sucrose | 238.67 | 955 |
| Cellulose | 40 | |
| Inulin | 10 | |
| Soybean Oil | 25 | 225 |
| Lard | 75.6 | 680 |
| Butter | 50.4 | 454 |
| Mineral Mix S10026 | 10 | |
| Dicalcium Phosphate | 13 | |
| Calcium Carbonate | 5.5 | |
| Potassium Citrate Monohydrate | 16.5 | |
| Vitamin Mix V10001 | 10 | 40 |
| Choline Bitartrate | 2 | |
| Cholesterol | 0.2943 | |
| FD&C Yellow Dye #5 | 0.025 | |
| FD&C Red Dye #40 | 0.025 | |
| **Total** | 922.8243 | 4057 |

**Table S3. Primer sequences for reverse transcriptase qPCR.**

| Gene Symbol | FORWARD | REVERSE | Annealing Temp. °C |
|---|---|---|---|
| *R18s* | GTAACCCGTTGAACCCCATT | CCATCCAATCGGTAGTAGCG | 55 |
| *B2m* | CTGCTACGTAACACAGTTCCACCC | CATGATGCTTGATCACATGTCTCG | 55 |
| *Gapdh* | ACAGTCCATGCCATCACTGCC | GCCTGCTTCACCACCTTCTTG | 55 |
| *Acadm* | ATTGCCAATCAGCTAGCCAC | CTGATAGATCTTGGCGTCCC | 56 |
| *Acsl5* | AATGTGTTCAAAGGCTACCTAAAGGACCC | GCGACCAATGTCCCCAGTGTGA | 62 |
| *Acox* | AGCGAGCCAGAGCCCCAG | TCAGGCAGCTCACTCAGG | 59 |
| *Adipoq* | GCACTGGCAAGTTCTACTGCAA | GTAGGTGAAGAGAACGGCCTTGT | 55 |
| *Apob* | CAGGTGGCCACAGCCAATAA | ACTGCAGGTCTGGCTCAGGA | 61 |
| *Atgl* | GACCTGATGACCACCCTTTC | TGCTACCCGTCTGCTCTTTC | 57 |
| *Cd36* | GGTGATGTTTGTTGCTTTTATGATTTC | TGTAGATCGGCTTTACCAAAGATG | 54 |
| *Dgat1* | CCCATACCCGGGACAAAGAC | ATCAGCATCACCACACACCA | 59 |
| *Fasn* | GCTGCGGAAACTTCAGGAAAT | AGAGACGTGTCACTCCTGGACTT | 58 |
| *Lep* | GCTACAGGCCTTTTGTTGGC | CCACAGAATGGGTGGGAGAC | 55 |
| *Lpin1* | AAGAGACTGACAACGATCAGGA | TTCCCCAGAGAACCAGTGGAT | 57 |
| *Lpl* | ATGGATGGACGGTAACGGGAA | CCCGATACAACCAGTCTACTACA | 58 |
| *Mogat1* | TGGTGCCAGTTTGGTTCCAG | TGCTCTGAGGTCGGGTTCA | 61 |
| *Mttp* | CTCTTGGCAGTGCTTTTTCTCT | GAGCTTGTATAGCCGCTCATT | 56 |
| *Pgc1a* | AACAAGCACTTCGGTCATCCCTG | TTACTGAAGTCGCCATCCCTTAG | 55 |
| *Plin1* | GGGACCTGTGAGTGCTTCC | GTATTGAAGAGCCGGGATCTTTT | 55 |

**Table S4. Statistical analyses of fold-change in liver triacylglyceride classes**

Attached in a separate Excel file.

**Table S5. ANOVA of liver triacylglycerides**

Attached in a separate Excel file.

**Table S6. Statistical analyses of fold-change in serum triacylglyceride classes**

Attached in a separate Excel file.

**Table S7. ANOVA of serum triacylglycerides**

Attached in a separate Excel file.

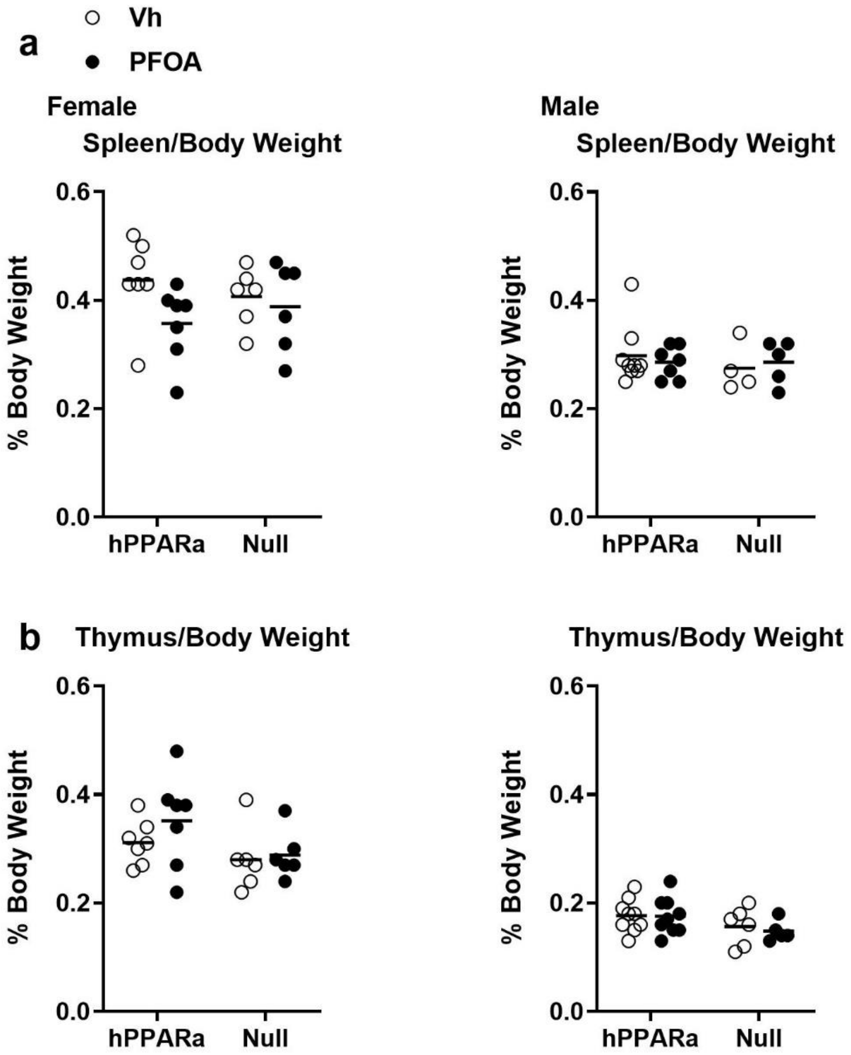

**Figure S1. Spleen/body weight (A) and thymus/body weight (B) ratios in PFOA-treated mice.** Three-week-old male and female humanized PPARα and KO mice were treated with either vehicle (Vh, NERL water with 5% sucrose) or PFOA (8 µM in NERL water with 5% sucrose) as drinking water for 6 weeks. During treatment, the mice were fed an American Diet (see Table S1). Thymus, spleen and body weights were determined at euthanasia. Data are from individual mice, with the mean indicated by a line. N = 5-9. There are no statistically significant differences (2 Factor ANOVA).

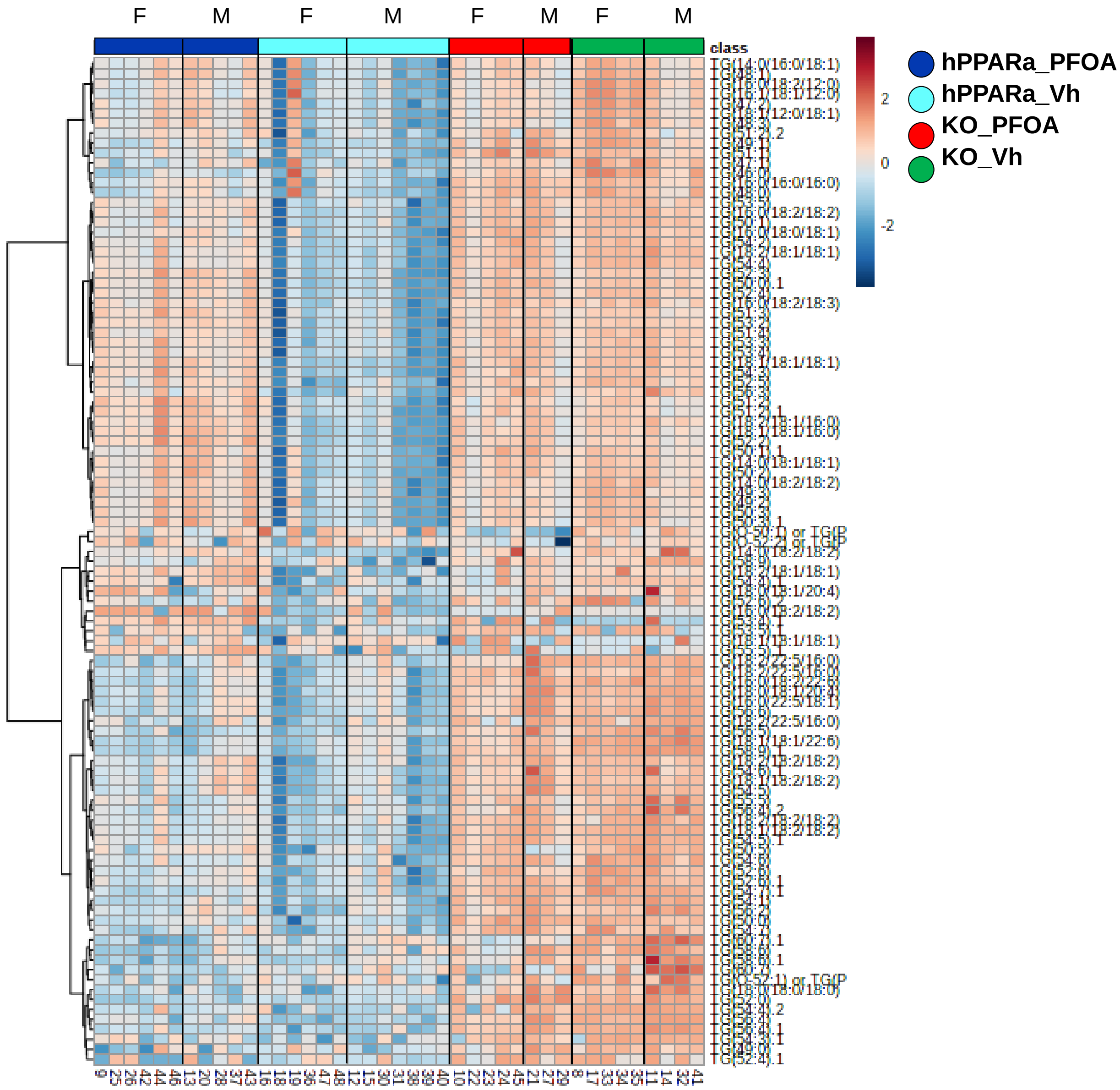

**Figure S2. Triacylglyceride profiles of liver.**
hPPARα and PPARα null mice were treated with vehicle or PFOA in drinking water for 6 weeks, as described in Fig. S1. TGs were identified and quantified by UHPLC-QTOFMS. N = 3-6.

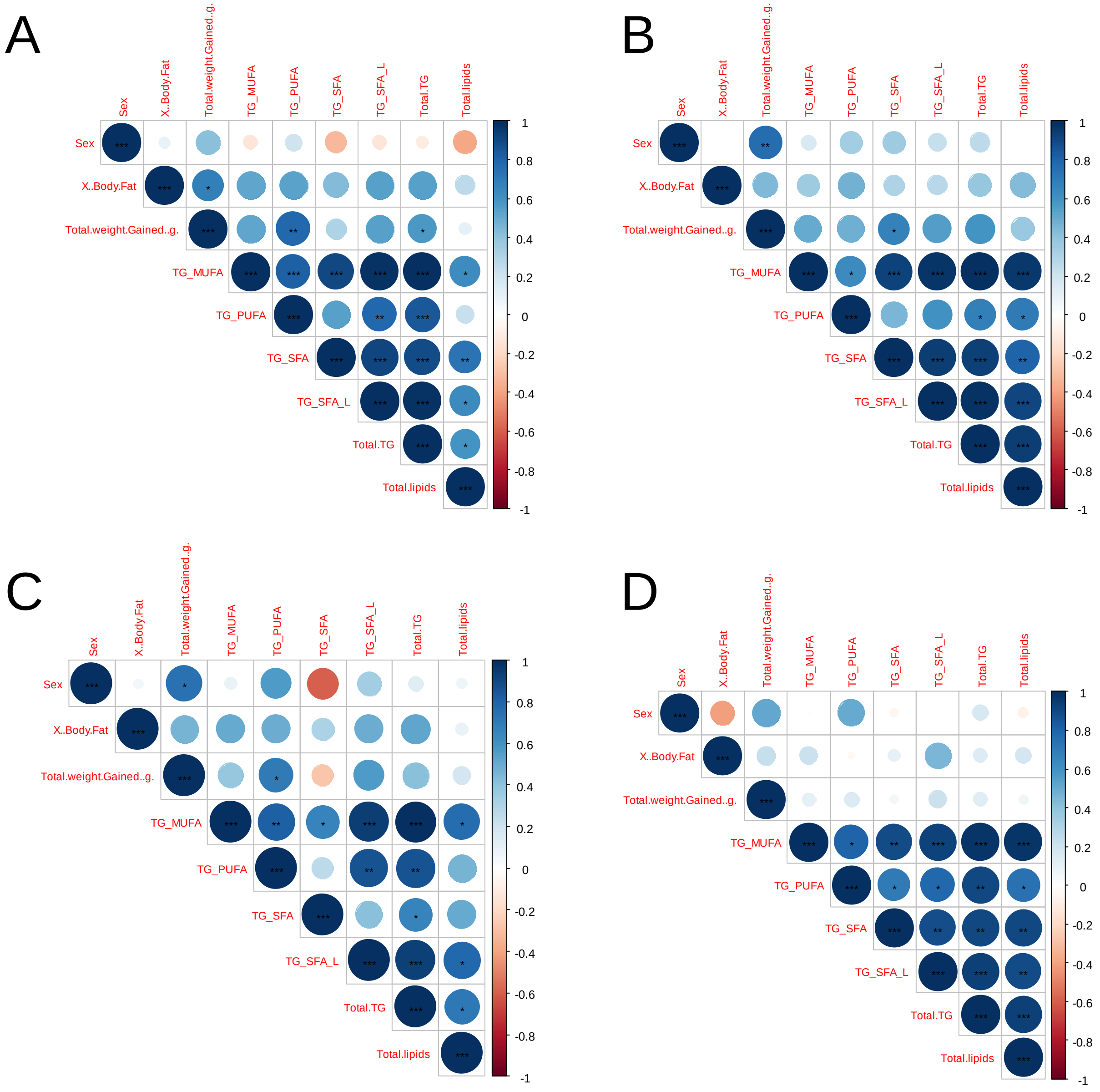

**Figure S3. Spearman correlation analysis of liver TG classes.**
hPPARα and PPARα null mice were treated with vehicle or PFOA in drinking water for 6 weeks, as described in Fig. S1. TGs were identified and quantified by UHPLC-QTOFMS. N = 3-6. A) hPPARα _Vh, B) hPPARα _PFOA, C) hPPARα null_Vh and D) hPPARα null_PFOA.
